# Decreasing alertness modulates perceptual decision-making

**DOI:** 10.1101/2020.07.23.218727

**Authors:** Sridhar R. Jagannathan, Corinne A. Bareham, Tristan A. Bekinschtein

**Affiliations:** Department of Psychology, University of Cambridge, Cambridge, United Kingdom; Department of Clinical Neurosciences, University of Cambridge, Cambridge, United Kingdom; School of Psychology, Victoria University of Wellington, Wellington, New Zealand; Institute of Neurophysiology, Charité Universitätsmedizin Berlin, Berlin, Germany; School of Psychology, Massey University, Palmerston North, New Zealand

## Abstract

The ability to make decisions based on external information, prior knowledge and evidence, is a crucial aspect of cognition and may determine the success and survival of an organism. Despite extensive work on decision-making mechanisms/models, understanding the effects of alertness on neural and cognitive processes remain limited. Here we use electroencephalography and behavioural modelling to characterise cognitive and neural dynamics of perceptual decision-making in awake/low alertness periods in humans (14 male, 18 female) and characterise the compensatory mechanisms as alertness decreases. Well-rested human participants, changing between full-wakefulness and low alertness, performed an auditory tone-localisation task and its behavioural dynamics was quantified with psychophysics, signal detection theory and drift-diffusion modelling, revealing slower reaction times, inattention to the left side of space, and a lower rate of evidence accumulation in periods of low alertness. Unconstrained multivariate pattern analysis (decoding) showed a ~280ms delayed onset driven by low alertness of the neural signatures differentiating between left and right decision, with a spatial reconfiguration from centro-parietal to lateral frontal regions 150-360ms. To understand the neural compensatory mechanisms with decreasing alertness, we connected the evidence-accumulation behavioural parameter to the neural activity, showing in the early periods (125-325ms) a shift in the associated patterns from right parietal regions in awake, to right fronto-parietal during low alertness. This change in the neurobehavioural dynamics for central accumulation related cognitive processes define a clear reconfiguration of the brain networks’ regions and dynamics needed for the implementation of decision-making, revealing mechanisms of resilience of cognition when challenged by decreased alertness.

**Significance statement:** Most living organisms make multiple daily decisions and these require a degree of evidence from both the environment and the internal milieu. Such decisions are usually studied under sequential sampling models and involve making a behavioural choice based on sensory encoding, central accumulation, and motor implementation processes. Since there is little research on how decreasing alertness affects such cognitive processes, this study has looked at the cognitive and neural dynamics of perceptual decision-making in people while fully awake and in drowsy periods. Using computational modelling of behaviour and neural dynamics on human participants performing an auditory tone-localisation task, we reveal how low alertness modulates evidence accumulation related processes and its corresponding compensatory neural signatures.

## Introduction

The question of how decisions are made has shaped the world’s systems of government, justice and social order (Buchanan and O’Connell 2006). Studies on how the brain implements simple decisions have revealed several neurocognitive processes at the perceptual, central integration and motor implementation levels (Sigman and Dehaene 2005; O’Connell et al. 2018). However the modulatory effects of the internal milieu, homeostasis, alertness, and circadian influences on such processes have received less attention (Hull, Wright, and Czeisler 2003; Knowles 1993). Specifically, the effect of low alertness has only been tackled by sleep deprivation and brain injury studies but hardly by normal variations of wakefulness (Goupil and Bekinschtein 2012).

Perceptual decision-making in cognitive sciences has been successfully studied (Link and Heath 1975; Gold and Shadlen 2007) under sequential sampling models (SSMs), and consists of: a) Perceptual stage: a sensory system that transforms physical stimulus intensities to deliver decision-information; b) Central integration stage: a decision system that integrates and accumulates such decision-information variables and makes an optimal choice based on relative evidence; and c) Motor stage: a motor system that implements the appropriate motor plan/action. According to SSMs, accurate perceptual decisions separate the noise from the signal by repeatedly sampling and integrating evidence until there is enough in favour of one of the decision choices. The preferred concept to understand and develop a hypothesis is called the decision variable, and it is an accumulation of priors, evidence, internal milieu and value into a quantity that is interpreted by the decision rule to produce a choice (Gold and Shadlen 2007). This study aims to investigate how alertness modulates such decision mechanisms in spatial auditory perception.

To understand how alertness modulates perceptual decision-making, we need to first define the specific aspect of wakefulness to be used as the experimental manipulation (T. Bekinschtein et al. 2009). Changes in alertness can be classified into ‘tonic’ that span multiple trials/time-periods, and ‘phasic’ moment to moment changes produced in response to an ongoing task. A few recent studies have shown that alertness, measured by brain-stem systems indirect markers (pupil response), modulates individual decision making in moment to moment fluctuations (phasic) (McGinley et al. 2015; de Gee et al. 2017). Further to this, van Kempen and collaborators (van Kempen et al. 2019) showed that lower tonic and higher phasic alertness via pupil measurements, predicted shorter reaction times and was associated with a centroparietal positivity in EEG space. However, these studies have used pupil responses which in neural terms is an indirect marker for alertness, and can also be influenced by stimulus features like visual contrast and several other parameters (Wang et al. 2018).

Studies in stroke patients have also revealed the effect of alertness on cognitive processes like spatial attention (Robertson et al. 1998), in particular, lesions on one side of the brain create a difficulty in localising and paying attention to information on the side opposite to the lesion, a condition referred to as unilateral spatial neglect and usually more persistent after a stroke in the right hemisphere (Karnath and Zihl 2003). We have also shown a while back, mimicking the behaviour of hemispatial neglect patients (Langner and Eickhoff 2013), that during drowsy periods, even normal participants show more errors to the left side of the space in an auditory task (Bareham et al. 2015). However, these alertness related effects have not been studied in a perceptual decision-making framework which further motivated this study. Hence, we took the opportunity to merge the questions on spatial attention and perceptual decision-making modulated by alertness by implementing an auditory spatial attention task that we developed previously. This paradigm is well defined for this alertness modulation design, and robust and systematic results have been replicated with it. Further, the auditory stimuli can be delivered with eyes closed, and finally it has a simple response option which is appropriate for drowsy participants.

Thus, we designed a study to directly measure tonic alertness and understand its effect on decision-making. Here, we also utilised a recently developed computational method (Jagannathan et al. 2018) to measure alertness directly from EEG, and in combination with multi-level modelling and psychophysics, to understand its effect on behaviour. Next, we used a drift-diffusion model to parametrise the different elements of the decision-making process, neural decoding for data-driven characterization, and finally, connected the drift-diffusion model to the neural markers to reveal the compensatory mechanisms of decision-making and spatial attention.

## Materials and Methods

### Participants

Forty-one healthy human participants (no auditory, neurological or psychiatric abnormalities) were recruited. Data from 8 participants was discarded due to technical problems with headphone amplifiers (battery issue was only discovered post-hoc) and 1 participant for not following task instructions (switching response hand half way through the experiment). Thus only data from 32 participants (14 males, age: 24.46 ± 3.72 years old) was considered for further analysis. All participants were self-reported to be right-handed. This was also established by using Edinburgh handedness scale (Oldfield 1971) and each participant had a score of above 0 (right-handed) with mean 80.26 ± 23.59. Only easy sleepers (as per self report) were recruited and further they were administered with the Epworth sleepiness scale (Johns 1991) on the day of the experiment. 29 participants had a sleepiness score >= 7 (classified as easy sleepers) and 3 of them had a sleepiness score >=4. All participants were asked not to consume any stimulant like coffee/tea before the experiment that day. The study and the experimental protocol was approved by the cambridge psychology research ethics committee and written informed consent was provided by all participants. A monetary compensation of £30 was provided for participation.

### Experimental task

Each participant underwent two experimental sessions a) Alert b) Drowsy. The alert session lasted approximately 8 minutes in duration, with the participants seated upright and lights on. Further they were instructed to stay awake throughout this session. Followed by which the drowsy session was done, which lasted approximately 1.5-2 hours in duration, with the participants reclined to maximum in a chair and lights off to promote drowsiness. Further they were provided with a pillow for neck support and were allowed to fall asleep. It is critical to note that the inter-trial interval in the alert session is 2-3 sec, whereas in the drowsy session it is 4-5 sec. This increase in intertrial interval, longer duration of session was intended to promote drowsiness in the drowsy session (Kosslyn and Andersen 1995; Bareham et al. 2014). Before the start of the experiment, the participants were allowed a practise session to become familiar with the task. The trial details for individual session are given below:

### Alert session

Participants were presented with 124 complex harmonic tones (guitar chords) that fell on the left or right side of their veridical midline (0°) ranging from −59.31° to +59.31°. These tones were recorded using in-ear microphones in free-field (Bareham et al. 2014). Six tones from −59.31° to −39.26° were presented two times each; twelve tones from −35.24° to - 1.86° were presented four times each. A similar pattern was repeated on the right side with twelve tones from 1.86° to 35.24° presented four times each, six tones from 39.26° to 59.31° presented two times each. The tones in the midline (0°) were presented four times, resulting in a total of 124 tones. The order of tones presented was randomized per participant. Further, participants were instructed to keep their eyes closed and respond (as quickly and as accurately as possible) with a button press (using left/right thumb) indicating the location of the tone (left or right). Each trial began after a random interval of 2-3 seconds and if the participant did not respond for 5 seconds, the next trial was started.

### Drowsy session

Participants were presented with 740 complex harmonic tones (as above) that fell on the left or right of their veridical midline (0°) again ranging from −59.31° to +59.31°. Six tones from −59.31° to −39.26° were presented twenty times each; twelve tones from −35.24° to −1.86° were presented twenty times each. Similar pattern was repeated on the right side with twelve tones from 1.86° to 35.24° being presented twenty times each, six tones from 39.26° to 59.31° presented twenty times each. The tone in the midline (0°) was presented twenty times, resulting in a total of 740 tones. The order of tones was again randomized per participant. Participants were again instructed to keep their eyes closed and respond (as quickly and as accurately as possible) with a button press (by left/right thumb) indicating the direction of the tone (left or right). Each trial began after a random interval of 4-5 seconds and if the participant did not respond for 5 seconds, the next trial was started. The participants were gently awoken if they didn’t respond to more than 3 trials consecutively.

### EEG recordings and preprocessing

EEG data was acquired with 128 Ag/AgCl electrodes (Electrical Geodesics Inc) using Cz as the reference electrode. The impedances of all electrodes were kept below 100 KΩ (to ensure higher signal to noise ratio) and data was acquired at a sampling rate of 500 Hz. EEG data was pre-processed with custom made scripts in MATLAB (MathWorks Inc. Natick, MA, USA) using EEGLAB toolbox (Delorme and Makeig 2004). The preprocessing steps are as follows: First, the peripheral electrodes that covered the regions of forehead, cheeks and neck were removed to reduce artifacts related to eye and muscle movements, thus retaining only 92 channels that covered the scalp. Second, the data was bandpass filtered with zero phase shift between 1 and 40 Hz using hamming windowed-sinc FIR filter and further resampled to 250 Hz. Third, pre-trial and post-trial epochs per trial were created. For the pre-trial epochs, the data was epoched from −4000ms to 0ms prior to the onset of the stimuli. The pre-trial epochs were created only in the drowsy session and not in the alert session (details below). For the post-trial epochs, the data was epoched from −200ms to 800ms to the onset of the stimuli for both the alert and drowsy sessions. Fourth, the trials that exceeded the amplitude threshold of ±250 μV were removed in a semi-automatic fashion. Fifth, the bad channels were detected in a two-step fashion: a) channels are considered bad (zero activity) if channel variance is below 0.5. b) The normalized power spectrum of the remaining channels was computed and any channel that exceeded the mean power spectrum by ±3 standard deviations was marked bad. Sixth, to remove further artifacts related to eye-blink and muscle movement, independent component analysis (ICA) was performed on the channels not marked as bad in the previous step. ICA components that correspond to artifacts were rejected by manual inspection. Seventh, the bad channels were now interpolated using spherical interpolation. Eighth, the bad trials were detected again using an amplitude threshold of ±250 μV and bad electrodes (those exceeding the threshold) in such trials were interpolated in a trial-by-trial fashion. Ninth, the post-trial epochs were re-referenced to the average of all channels (whereas the pre-trial epochs were maintained with the recorded reference - Cz).

### Alertness level classification

The preliminary step in both behavioural and neural analysis is to classify trials into ‘alert’ and ‘drowsy’. The data from the pre-trial epochs were used to classify each trial into alert or drowsy. For the alert session, all pre-trial periods were considered to be ‘alert’ as participants were explicitly instructed to stay awake with short inter-trial intervals of 2 sec and overall shorter duration of session (both promote wakefulness) and lights switched on, with upright seating. Consequently, none of the participants failed to respond to any of the trials in the alert session which supported our assumption. In the drowsy session, the participants were allowed to fall asleep with longer inter-trial intervals of 4-5 sec and lights switched off, with seating reclined to maximum with a pillow for the head support and hence several participants failed to respond to some trials in the drowsy session. For each trial in the drowsy session, pre-trial epochs were analysed using the micro-measures algorithm (Jagannathan et al. 2018). Briefly, the micro-measures algorithm operates on 4-sec epochs in a two-step fashion. In the first step, ‘alert(relaxed)’ trials are separated from the ‘drowsy’ trials (sub-divided into ‘mild’ and ‘severe’) by using a combination of features like: variance explained by different frequency band predictors and coherence at different frequency bands (See (Jagannathan et al. 2018) for details). For predictor variance the different features were computed by first generating predictors based on spectral variation in different frequency bands A: 2–4 Hz; B: 8–10 Hz, C: 10–12 Hz; D: 2–6 Hz in the occipital electrodes (E75, E70, E83) and then fitting them per electrode per epoch. For coherence, the frequency bins: Delta:[1–4 Hz], Theta:[4–7 Hz], Alpha:[7–12 Hz], Sigma:[12–16 Hz], Gamma:[16–30 Hz] were used at the occipital: E75 (Oz), E70 (O1), E83 (O2), frontal: E33 (F7), E122 (F8), E11 (Fz), central: E36 (C3), E104 (C4), temporal: E45 (T3-T7), E108 (T4-T8), E102 (TP8), E115, E100 (TP10) sites. These electrode numbers can be identified using the GSN hydrocel 128 channel map from EGI and corresponding locations in the standard 10-10 system are given in brackets wherever identified (Luu and Ferree 2005). In the second step, ‘drowsy(severe)’ trails were further computed using a combination of grapho-element identification like vertex waves, spindles etc. We only used the ‘drowsy(mild)’ trials for identifying ‘drowsy’ trials, as the participants usually do not respond under ‘drowsy(severe)’ trials. We similarly excluded the ‘alert(relaxed)’ trials in the drowsy session to instead compare the drowsy trials of the drowsy session to all of the trials in the alert session. It should be noted that we could not use the micro-measure algorithm on the alert session as mainly the pre-trial duration was only 2-sec which precludes its application (as 4-sec is the recommended for the algorithm). However, none of the participants failed to respond in the alert session as mentioned before. Finally to cross-validate our assumptions, we performed a t-test on the distributions of reaction times of the ‘alert’ versus the ‘drowsy’ trials per participant. 28 of the 32 participants significantly differed in reaction time distributions which is a direct effect of alertness (Ogilvie 2001), as participants tend to be slower and produce more variable reaction times, when drowsy.

The details of this analysis are present here: https://github.com/SridharJagannathan/decAlertnessDecisionmaking_JNeuroscience2021/blob/main/Scripts/notebooks/Figure1supplement_RT_persubject.ipynb

### Behavioural analysis

#### Error proportion

In order to understand how the rate of errors differs across different stimuli (left or right tones) and how it is modulated by alertness levels (alert or drowsy) we performed the following analysis. First, we computed the proportion of errors made by each participant under each alertness level (alert, drowsy) and under each stimulus type (left or right tone). If the total number of trials for any participant under any condition is less than 5, then the corresponding error proportion (for that condition) is ignored in the analysis. We decided to use multilevel models for this analysis over traditional repeated measures of analysis of variance, because different participants had different levels of alertness (specifically differing number of alert, drowsy trials per condition). We defined 4 different multilevel models to understand the modulation of error proportion by state of participant (alert, drowsy) and stimulus (left, right). In the null model, the error proportion depends only on the mean per participant (fixed effect) and the participant id (random effect). In the second model (state model), the error proportion depends only on the state of the participant (fixed effect) and the participant id (random effect). In the third model (stimulus model), the error proportion depends only on the stimulus (fixed effect) and the participant id (random effect). In the fourth model (state-stimulus model), the error proportion depends on a combination of the state of the participant and the stimulus, both used as fixed effects and the participant id (random effect). These 4 models were fitted using the ‘lmer’ function (‘lmerTest’ package) in R (Kuznetsova, Brockhoff, and Christensen 2017) and the winning model is identified as the one with the highest log-likelihood by comparing it with the null model and performing a likelihood ratio chi-square test (χ^2^). Finally the top two winning models were compared against each other using ‘anova’ function in R (Fox and Weisberg 2018), to validate whether the winning model (if it is more complex) is actually better than the losing model (if it is simpler). The state-stimulus model emerged as the winning model.

The details of this analysis are present here: https://github.com/SridharJagannathan/decAlertnessDecisionmaking_JNeuroscience2021/blob/main/Scripts/notebooks/Figure2a_errorproportion.ipynb

The different models along with their log-likelihood values are shown below:

**Table 1:**
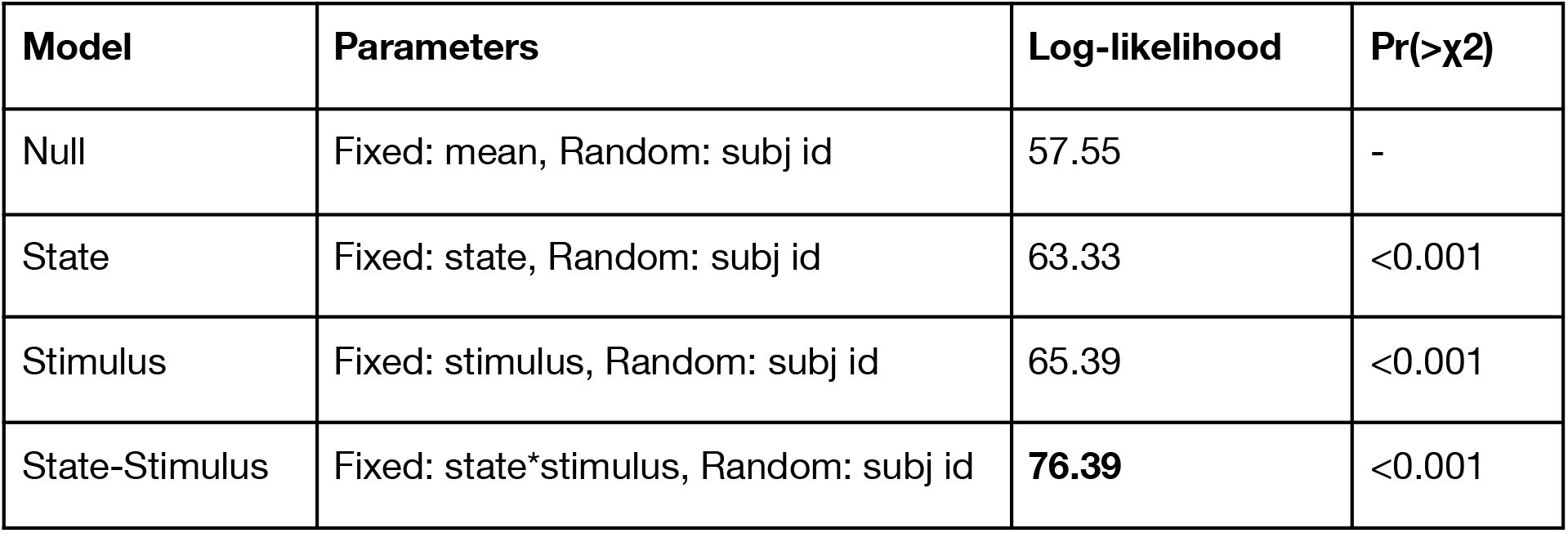
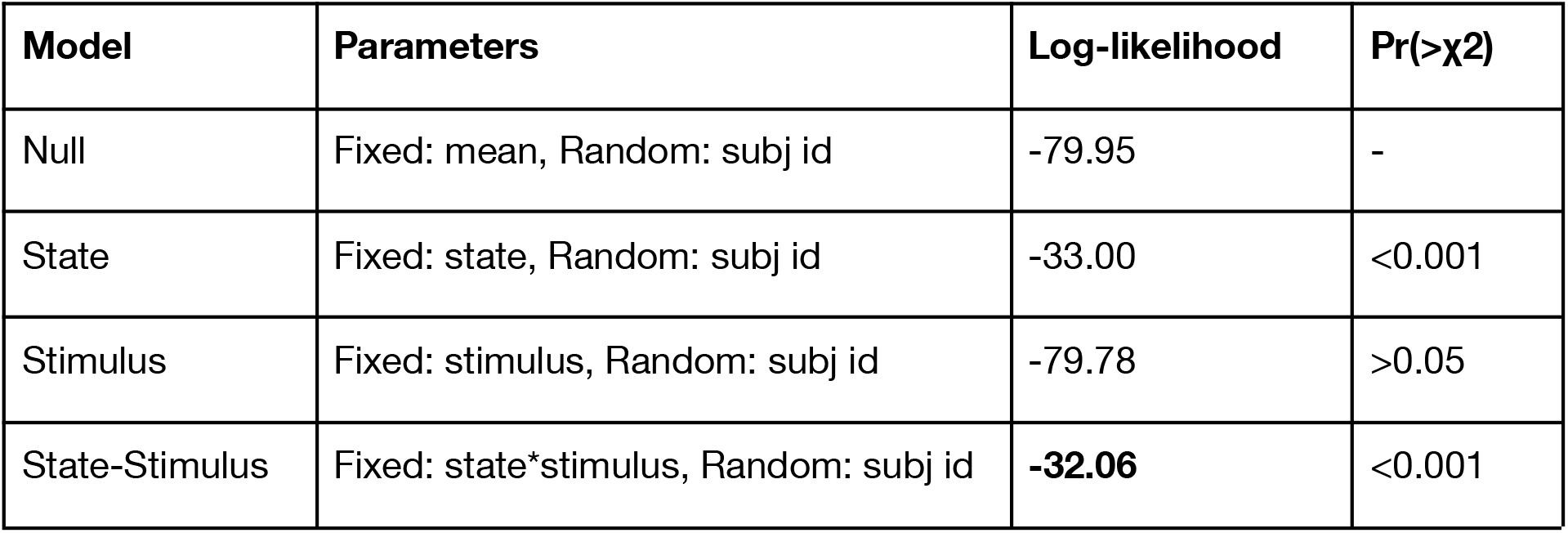
Model comparison

**Table 2:**
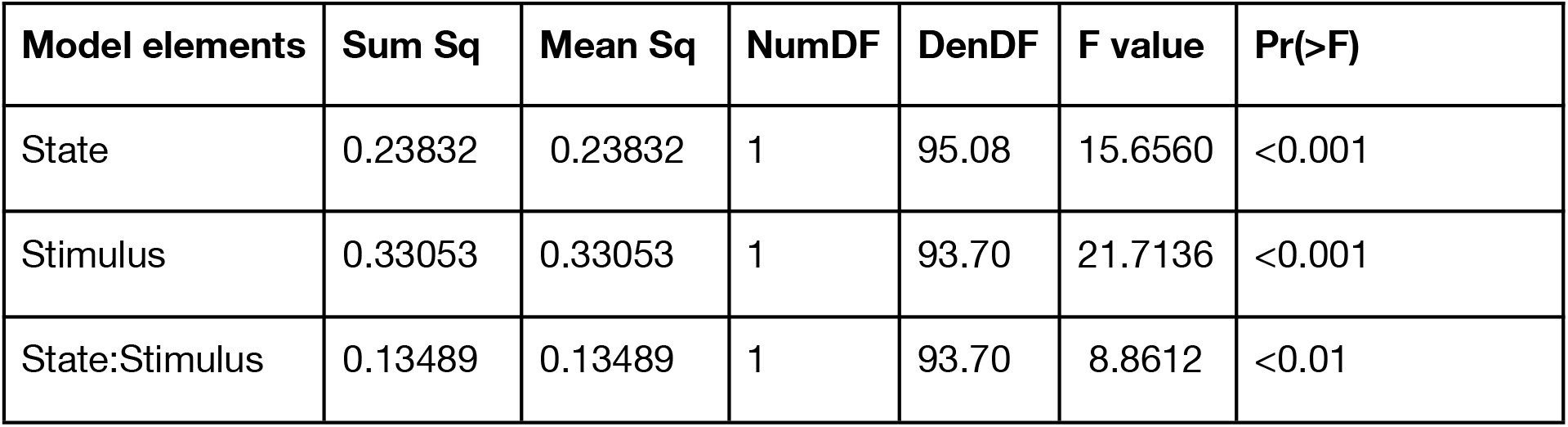
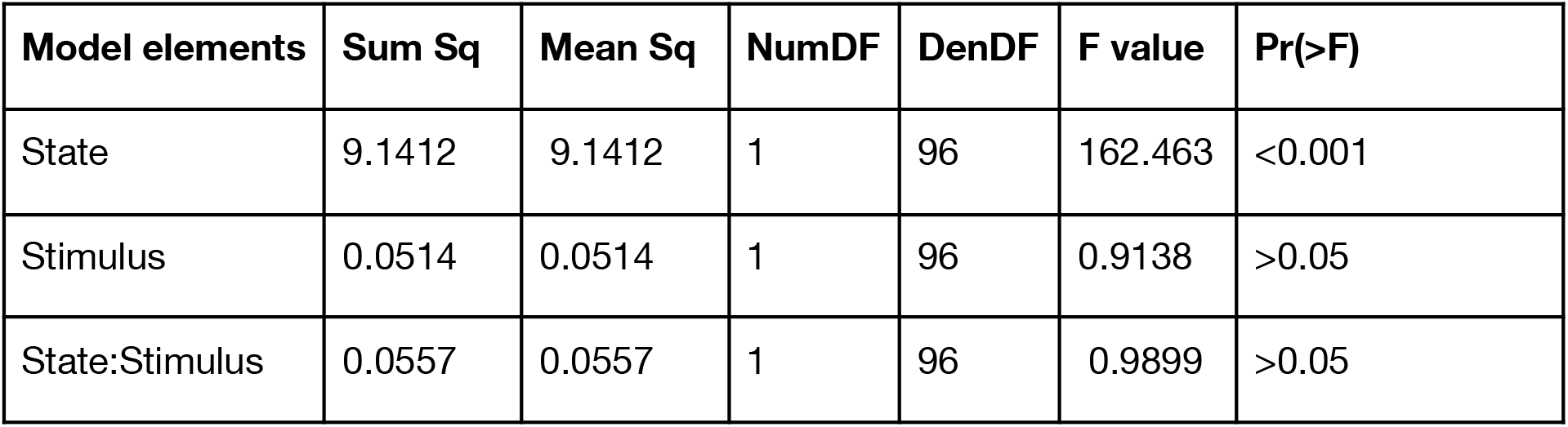
Type III analysis of variance with Satterthwaite’s method of the winning model (State-Stimulus)

The state-stimulus (winning model) was further analysed with the ‘anova’ function and it was found that there was a reliable main effect of both state (p<0.001) and stimulus (p<0.001) on error proportion and also the interaction between state and stimulus also had a reliable effect (p<0.01) on error proportion. Further, we performed post-hoc tests using tukey adjustment (for multiple comparisons) to identify differences between pairs that are significant.

### Reaction times

In order to understand how the reaction times differ across different stimuli (left or right tones) and how it is modulated by alertness levels (alert or drowsy) we performed a multilevel modelling analysis similar to the analysis on error proportion as above. First, we computed the reaction times of each participant under each alertness level (alert, drowsy) and under each stimulus type (left or right tone). We defined 4 different multilevel models to understand the modulation of reaction times by state of participant (alert, drowsy) and stimulus (left, right). In the null model, the reaction time depends only on the mean per participant (fixed effect) and the participant id (random effect). In the second model (state model), the reaction time depends only on the state of the participant (fixed effect) and the participant id (random effect). In the third model (stimulus model), the reaction time depends only on the stimulus (fixed effect) and the participant id (random effect). In the fourth model (state-stimulus model), the reaction time depends on a combination of the state of the participant and the stimulus, both used as fixed effects and the participant id (random effect). The state-stimulus model emerged as the winning model.

The details of this analysis are present here: https://github.com/SridharJagannathan/decAlertnessDecisionmaking_JNeuroscience2021/blob/main/Scripts/notebooks/Figure2supplement_RT.ipynb

The different models along with their log-likelihood values are shown below:

The state-stimulus (winning model) was further analysed with the ‘anova’ function and it was found that there was a reliable main effect of only state (p<0.001) on reaction time.

### Subjective midline shifts

The change in subjective midline was quantified by fitting psychometric functions to the responses produced by each participant under alert and drowsy conditions. The proportion of rightward responses for each participant under each stimulus condition from −60° to +60° was fitted with a cumulative normal function using the ‘quickpsy’ (Linares and López-Moliner 2016) package in R. In order to evaluate the different parameters involved in the modulation of psychometric functions, we evaluated two different models. The first model was where only the mean of the function shifted for individual participants across alert and drowsy conditions. The second model was where both the mean and the slope shifted across conditions. Model selection was performed using the Akaike Information Criterion, model fits from 9 subjects favoured the first model and 23 subjects favoured the second model. Thus we used the model where both the mean and the slope varied across alert and drowsy conditions. The mean of the cumulative normal function (the point where the curve crosses 0.5 in the y-axis) is also referred to as the subjective midline (‘bias’). The subjective midline is the stimulus (angle) at which the participant performs at chance (0.5), which in an ideal world would be closer to the veridical midline (0°). The slope of the cumulative function represents the ‘sensitivity’ of the system. In general, large variations in the bias point tend to reduce the sensitivity of the system. Further for performing a paired t-test on the change in the parameters across alert and drowsy conditions we also ignored 13 participants that had a bias point outside ±60° (as the overall stimulus angle can vary only between −60° to +60°) and standard deviation of more than 30°. The details of this analysis are present here: https://github.com/SridharJagannathan/decAlertnessDecisionmaking_JNeuroscience2021/blob/main/Scripts/notebooks/Figure2c_d_e_psychophysics_biasshift.ipynb

### Drift-diffusion modelling

The different elements of the decision-making process can be teased apart by using the drift-diffusion model. The parameters of this model include: drift-rate (v) -- rate of evidence accumulation, boundary separation distance (a) -- distance between the two decision boundaries, non-decision time (t) -- for accounting other processes like stimulus encoding (before the start of evidence accumulation process), response execution (after the end of evidence accumulation). Further the different sources of bias that can be modelled are: bias point (z) -- bias in the starting point or drift criterion (dc) - a constant factor (slope) added to the drift rate. We implemented the drift diffusion process using a hierarchical drift diffusion model (HDDM) (Wiecki, Sofer, and Frank 2013) that allows for a hierarchical Bayesian procedure to estimate the model parameters. The principal reason for using such hierarchical procedures is because different participants fall asleep in different ways (differing number of alert and drowsy trials), hence usage of hierarchical bayesian procedure allows for robust estimation of model parameters under such limited conditions of trials (Zhang et al. 2016). We used the response of each participant (‘left’ or ‘right’ button press) instead of accuracy (‘correct’ or ‘incorrect’) to fit the HDDM. Such a procedure is referred to as stimulus-coding and allows for robust estimation of sources of bias without being affected by accuracy. In the first step, we decided to identify the sources of bias (with z or dc). Several studies across auditory and visual modalities (Stelmach and Herdman 1991; Benwell, Harvey, and Thut 2014) have shown existence of an initial spatial bias which is modulated by different factors like time-on-task and alertness levels. Hence it is important to identify and parametrise the source of this bias to either response driven (changes in z) or stimulus driven (changes in dc) by using model comparison techniques (such as χ^2^ or based on information criterion) as done in previous studies (White and Poldrack 2014; Leite and Ratcliff 2011). For this we implemented 8 different models that allowed the parameters (z,v) to vary depending on state (alert or drowsy) or stimulus (left or right). Similarly we implemented 8 different models that allowed the parameters (dc,v) to vary depending on state or stimulus. In each model, 15000 samples from the posterior distribution were estimated using Markov chain Monte Carlo methods. 5000 samples were further discarded as burn-in to discard the effect of initial values on the posterior distribution estimation. We then choose the best model among the bias models using the one with the lowest deviance information criterion (DIC). DIC allows for computing model accuracy while penalising for model complexity (Spiegelhalter et al. 2002). The best model among the bias models was where the v varied with state, stimulus and z varied with stimulus alone. In the next step, we used this best model and developed a set of combined models that allows variation in the parameters (a, t) with state or stimulus. We then choose the best model among the 4 different combined models. The final best model was one where ‘v’ varied with state, stimulus and ‘z’ varied with stimulus and ‘a’ varied with state and ‘t’ varied with stimulus alone. This final best model checked for model convergence using the Gelman-Rubin statistic. The convergence statistic was computed for 5 different runs and was found to be closer to 0.99 (values closer to 1 but less than 1.2 indicate convergence) (Gelman et al. 2013; Spiegelhalter et al. 2002).

The details of this analysis are present here: https://github.com/SridharJagannathan/decAlertnessDecisionmaking_JNeuroscience2021/blob/main/Scripts/notebooks/Figure3d_k_suppl_hddm.ipynb

### Neural analysis

#### Decoding

We used multivariate pattern analysis (MVPA) techniques to analyse the divergent patterns in EEG data. Specifically we used decoding in which patterns of brain activity are analysed in order to predict the experimental condition under which it was generated. Traditional ERP (event related potentials) analyses rely on using a-priori identified spatial locations or temporal segments in the data to measure the differences across conditions. However decoding techniques do not rely on a-priori definitions and perform much better in detecting differences across experimental conditions (Fahrenfort et al. 2018). Temporal decoding involves using EEG data (X) composed of size: [electrodes x time points x trials] to predict the stimuli presented (Y). The first step consists of fitting an estimator (w) to a subset of the data (X_train_) to predict a subset of the experimental condition (Y_train_). The second step involves using this trained estimator on another subset of the data (X_test_) to predict a subset of the experimental condition (Y_test_). The third step involves evaluating the performance of this estimator using a validation measure by comparing the prediction (Ŷ_test_) with the actual label (Y_test_).

### Estimator construction

First, the EEG data is subjected to a standard scaler (using StandardScaler() from scikit-learn) that removes the mean of the data and scales it by its variance. Second, we used logistic regression to estimate the model parameters for maximally separating the different experimental conditions. Third, we implemented temporal decoding by using the sliding estimator (using SlidingEstimator() from scikit-learn) to fit the logistic regression model per time-point.

### Cross validation

The EEG data was first downsampled to 100 Hz and further binary classification was performed between two conditions (left and right stimuli) separately across alert and drowsy conditions per individual participant. As the target of the classification was stimuli being presented we only considered the trials where the participant made the correct decision. Each participant was considered for classification only if they had at least 25 trials under each condition. Further 5-fold cross validation was performed such that 4 folds were used for training and the last fold was used as a testing set. The classifier performance was evaluated using Area Under the Curve (AUC) of the receiver-operating characteristic (ROC). It is implemented using ‘roc_auc’ in the sliding estimator function in scikit-learn. When AUC is about 0.5 the classifier performs at chance, while the AUC score of 1 has a very good separability across classes. We computed the AUC-ROC score per participant as the average of the score across all the cross-validation folds. Further, we smoothed the scores using a 3 point moving average to smooth out spurious fluctuations. In order to identify the reliability of the AUC score at the group level, we performed a cluster permutation test (participants x timepoints) using MNE (spatio_temporal_cluster_1samp_test) (Gramfort et al. 2013). Thus producing p-values per time point at the group level, from which time points where we can infer those regions where AUC is reliably different from chance (0.5) at the group level.

### Coefficients of patterns

The parameters of the decoding (performed above) are not neurophysiologically interpretable in a straightforward way (Haufe et al. 2014). Hence the parameters of the backward model (decoding) need to be transformed to produce the forward model. This is done by obtaining the coefficients of the estimator model per participant using the ‘get_coef’ function from MNE (‘_patterns’). For performing group statistics in electrode space, we used the same cluster permutation based approach as described earlier.

### Decoding of responses

To tease apart the process related to evidence accumulation from motor implementation, we decided to decode the response hand of the participant. The EEG data for the response decoding was created by epoching the data from −800ms to 200ms prior to the onset of the response from the participant. The response-locked trials were preprocessed in a similar fashion to the stimulus-locked trials. The alertness level for the corresponding response-locked trial was obtained from the labels of the stimulus-locked trial. The decoding and other methods used are similar to the stimulus-locked trials, except that the target of decoding is the response hand being pressed (left or right thumb).

The details of the stimulus decoding are present here: https://github.com/SridharJagannathan/decAlertnessDecisionmaking_JNeuroscience2021/blob/main/Scripts/notebooks/Figure4_5_temporaldecodingstimuli.ipynb

The details of the response decoding are present here: https://github.com/SridharJagannathan/decAlertnessDecisionmaking_JNeuroscience2021/blob/main/Scripts/notebooks/Figure6_temporaldecodingresponses.ipynb

### Neuro-behavioural analysis

#### Regression with drift-diffusion model

To identify the correlates of the evidence accumulation process, we used the model parameters generated by the drift diffusion model (combined best model). First, the ERP data (post trial epochs) were z-scored per electrode per trial. Second, the ERP data was baseline corrected with pre-trial data from −200ms to 0ms. Third, the ERP data was averaged every 50ms per electrode per trial to create 20 time points (−200ms to 800ms) per electrode per trial. Fourth, the ERP data were entered into a trial-by-trial regression with the drift rate using the HDDMRegressor from the HDDM toolbox (Wiecki, Sofer, and Frank 2013). The model parameters are allowed to vary as per the combined best model. We estimated the influence of ERP data on the drift-rate, as it was the only parameter shown to vary with respect to both the direction of stimulus and the alertness of the participants. Thus, we use drift parameter *v* as a dependent variable and regressed the same against ERP data as shown below

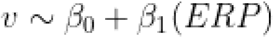

The above equation can be written in patsy form as below.

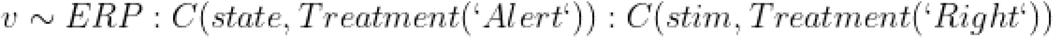

Here, the drift-rate covaries with the ERP value and depends on a combination of state with a reference value at ‘Alert’ and of stimulus with a reference value at ‘Right’.

The other model parameters are same as in the best model:

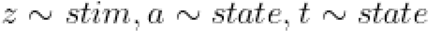

Here, *v* represents drift-rate, ERP represents z-scored ERP data per trial per time point per electrode, state represents alertness levels (‘alert’ or ‘drowsy’), stim represents stimulus types (‘left’ or ‘right’).

Fifth, the traces were computed per condition (state and stimuli combination). All the regressor models (1840 = 20*92 = timepoints*electrodes) for each participant were checked for convergence using the Gelman-Rubin statistic. The convergence statistic was computed for 3 different runs and was found to be closer to 0.99 (indicating convergence). Sixth, the differences in drift rate (between left and right stimuli) per time point per electrode were computed in both alert and drowsy condition. This difference represents the discriminability of the electrode in identifying the left and right stimuli at that time point. Thus this analysis yielded differences in drift rate per electrode, per time point, per condition, per participant. The differences can then be considered similar to the classifier patterns and can be analysed both in electrode and source spaces. Further we also computed group level differences using the cluster permutation techniques described earlier.

The details of this analysis are present here: https://github.com/SridharJagannathan/decAlertnessDecisionmaking_JNeuroscience2021/blob/main/Scripts/notebooks/Figure7_hddmregression.ipynb and comparison of patterns analysis is present here: https://github.com/SridharJagannathan/decAlertnessDecisionmaking_JNeuroscience2021/blob/main/Scripts/notebooks/Figure7_8_hddmregressionpatterns_comparison.ipynb

### Source reconstruction of the regression patterns

The coefficients created above can be projected into the source space to infer the brain regions involved in the pattern activity. Source reconstruction was achieved using a combination of Freesurfer (Fischl 2012) and MNE. First, the surface was reconstructed using ‘recon-all’ (Freesurfer) using the default ICBM152 template for the structural magnetic resonance image (MRIs). Next, the Boundary element model (BEM) was created using ‘make_watershed_bem’ (MNE). Further, scalp surfaces for the different element boundaries were created using ‘make_scalp_surface’ (MNE). Second, we performed the registration of the scalp surface with the default EEG channel locations (with fiducials as markers) manually using ‘coregistration’ (MNE). Third, the forward solution was computed using ‘make_bem_model’ (MNE). Fourth, to test if the source reconstruction of the electrode data is accurate we projected the ERP data of a sample participant into source space and analysed data from different regions of interest to confirm its validity. The patterns of each participant were then projected into source space in the following manner. First, we computed the noise covariance using the baseline data from −0.2 to 0 ms. Second, we used the forward solution and the noise covariance to create an inverse operator using ‘minimum_norm.make_inverse_operator’ from MNE. Third, used the individual pattern per participant and applied the inverse operator on it to produce the source reconstruction of the patterns per participant. For performing group statistics in the source space we used the same cluster permutation based approach (after moving them to the ‘fsaverage’ space) as described earlier.

The details of this analysis are present here: https://github.com/SridharJagannathan/decAlertnessDecisionmaking_JNeuroscience2021/blob/main/Scripts/notebooks/Figure9_hddmregressionpatterns_sources.ipynb

## Results

We organized the results from direct and model-free, to theoretically constrained. Error proportion, reaction times and subjective midline shifts are described as direct measures to evaluate the effects of alertness, while signal detection and drift-diffusion models theoretically constrain the interpretation of its parameters to perceptual, central and motor processes sequentially occurring during perceptual decision-making. We further organized the brain analyses similarly, using multivariate decoding to widely characterise the spatiotemporal neural signatures of the decision and, constrained by the modelled behavioural results, we map the neural dynamics of evidence accumulation in full wakefulness and low alertness. This approach allowed for both, open and exploratory neurobehavioural characterization and hypotheses-driven evaluation of the effects of low alertness on perceptual decision-making in spatial attention.

### Error-proportion modulated by alertness

First, we used multilevel modelling to understand how the errors made by each participant in the auditory tone-localization task was influenced by the stimulus presented (left or right tone) and the state of the participant (alert or drowsy). We defined 4 different multilevel (linear mixed) models where errors were modulated by various combinations of stimulus and alertness states (see methods section). The analysis of the variance table of the winning model shows that both alertness (F(1, 95.08) = 15.65, p<0.001) and stimulus type (F(1,93.70) = 21.71, p<0.001) have an effect on error-proportion. Further, there was a reliable interaction between alertness levels and stimulus type (F(1,93.70) = 8.86, p<0.01). Next, post-hoc analysis revealed a reliable difference between alert and drowsy conditions for left stimuli (p<0.001), and not for right stimuli (p = 0.899). These behavioural results (Figure 2A) replicate the findings of (Bareham et al. 2014) in an independent study, task design, alertness measure (and lab), reporting an increase in location assignment errors on tones from the left side of the midline when people became drowsy.

**Figure 1:**
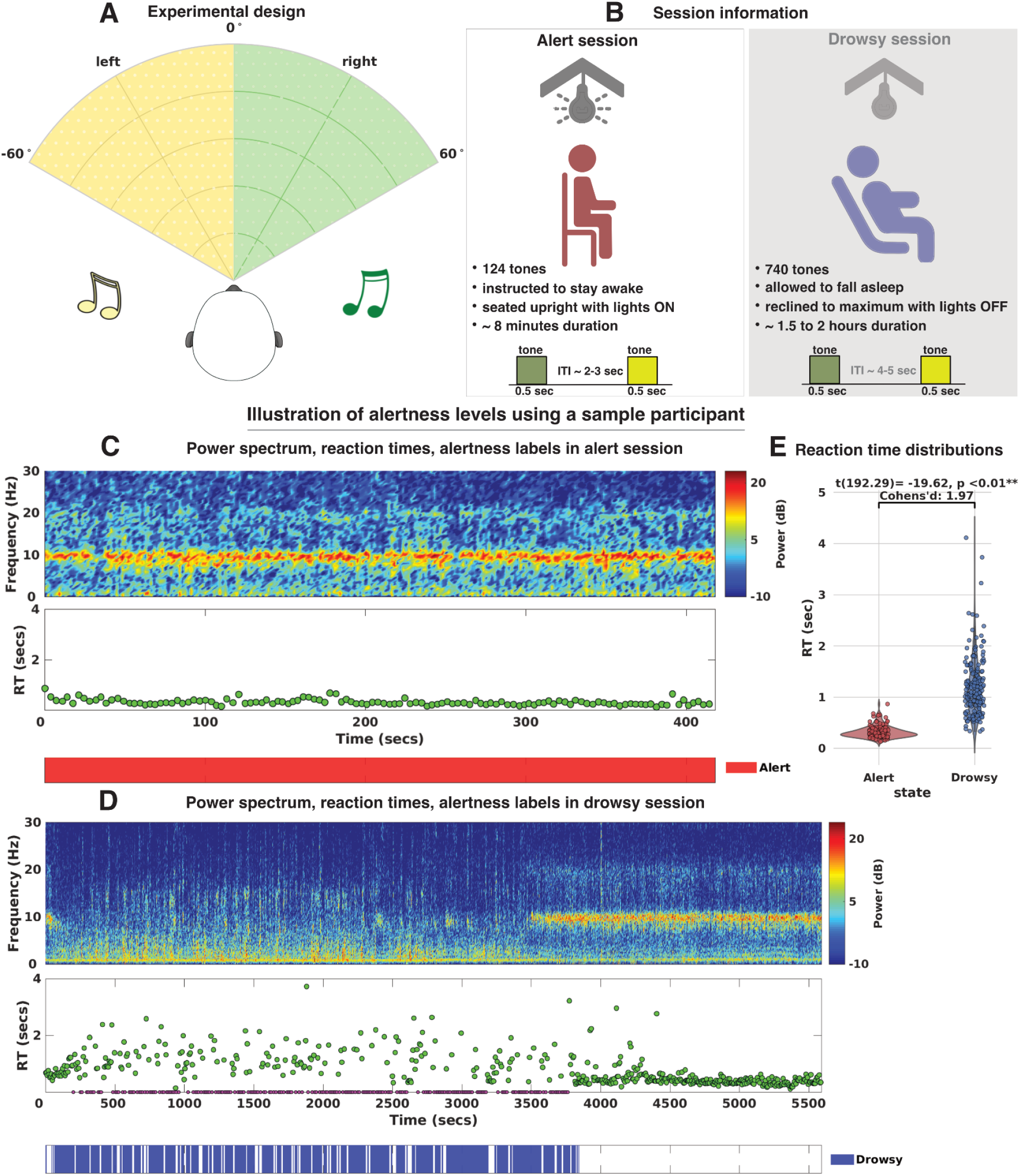
A) auditory spatial attention task: participants had to localize the direction of auditory tones coming from left and right side of the midline. B) session details: each participant underwent an alert and drowsy session. In the alert session of approximately 8 minutes in duration, participants are instructed to stay awake and seated upright with lights on. The shorter inter-trial interval of 2-3 seconds and shorter duration of the session ensured that none of the participants failed to respond in any of the trials. In the drowsy session of approximately 1.5 to 2 hours in duration, participants are allowed to fall asleep with the seat reclined to maximum with a pillow for head support. The longer inter-trial interval of 4-5 seconds and longer duration of the session ensured that most participants became drowsy. C,D) Alert and drowsy trial identification: All the trials in the alert session are considered to be alert (124), whereas among the 740 trials in the drowsy session, we employed the micro-measures algorithm (Jagannathan et al. 2018) and divided the pre-trial periods into different categories, among which we choose the mild drowsy trials. Here we can see that in the drowsy session this sample participant failed to respond several times (purple dots in RT plot) and the drowsy trial labels coincide in the nearby periods, whereas in the alert session the participant systematically responds. E) cross-validation of alert and drowsy trial labels per participant: for each participant we then performed a t-test comparing the distribution of reaction times of alert vs drowsy trials and found that for 28 of 32 participants, the distributions reliably differed, validating the approach (Jagannathan et al. 2018).

**Figure 2:**
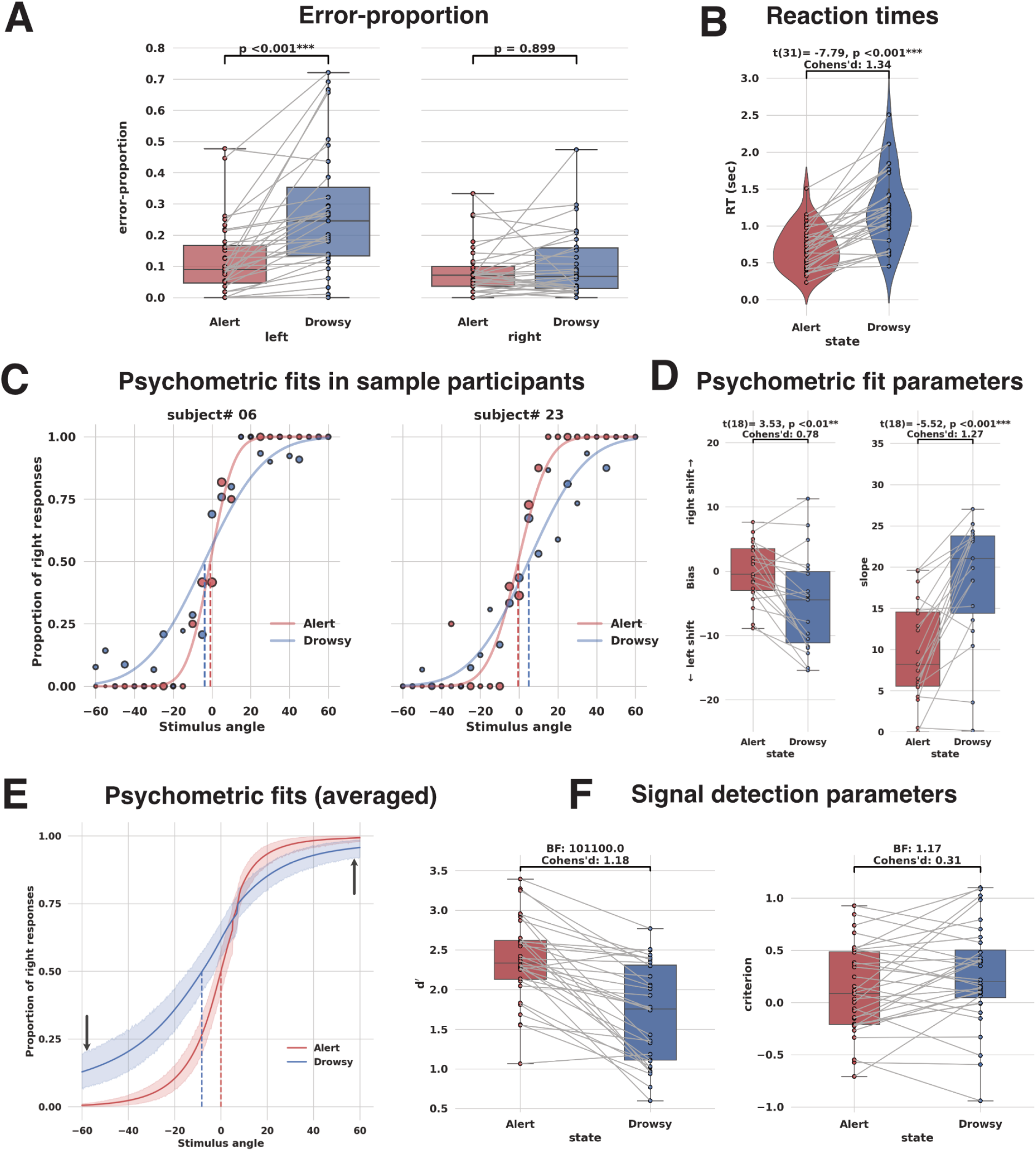
Behavioural analysis: A) proportion of errors committed in alert and drowsy periods across left and right stimuli. Multilevel modelling reveals that error rates depend on stimulus and state of the participant. Post-hoc tests indicate that error proportion is reliably modulated by alertness but only for left stimuli. *** indicates p<0.001, ns indicates not significant (or not reliable), error bars indicate standard error of the mean. B) Mean reaction times for individual participants. Reaction times are variable and slower under drowsy conditions and further a paired samples t-test indicates reliable difference (p<0.001) in reaction times across alert and drowsy periods. C) Example fits of two different participants are shown. For each participant under each condition (alert, drowsy), the proportion of rightward responses were fitted to the stimulus angle varying from −60° to +60°. The size of the dots represent the normalised (per condition) number of trials under each angle. Here, we notice that the bias point (dotted line) shifts towards the left side as the participant becomes drowsy. In other words, as the participant becomes drowsy they overestimate the right side of space. For another participant we notice the opposite effect. D) Spatial bias, Slope parameter per participant (from psychometric fits). Bias level shifts towards the left side (indicating more left errors) for most participants. Negative bias values indicate shifts in the subjective midline towards the left and positive values indicate shift towards the right. Further, a paired sample t-test indicates reliable difference (p<0.01). Slope values of the psychometric fits also increased as participants became drowsy, indicating shallower slopes. Further, a paired sample t-test indicates reliable difference (p<0.001). E) Group level psychometric fits indicate the shift in subjective midline (dotted lines), shaded regions are confidence interval bounds. Arrows indicate a gap in the asymptotes at −60° compared to +60° between alert and drowsy state, further evidence of inattention to the left side. F) Signal detection analysis shows that d′ (sensitivity) is strongly modulated by alertness in comparison to the c (criterion) using bayes factor.

### Reaction times are modulated by alertness

Second, we aimed to quantify the modulation in the response profiles (reaction times) of individual participants by alertness levels. A paired sample t-test with t(31) = −7.79, p<0.001 revealed a reliable effect of alertness on reaction time distributions. Further to this we also employed the framework of the multilevel modelling to understand how the reaction times of participants was influenced by the stimulus presented (left or right tone) and the state of the participant (alert or drowsy). We defined 4 different multilevel (linear mixed) models where errors were modulated by various combinations of stimulus and alertness states (see methods section). The analysis of the variance table of the winning model shows that only alertness (F(1, 96) = 162.46, p<0.001) has an effect on reaction times (convergent with the t-test before). These behavioural results (Figure 2B) converge with the original findings from Hori (Hori, Hayashi, and Morikawa 1994) indicating slower reaction times under lower levels of alertness, and in agreement to all our previous work (Noreika et al. 2020; Comsa, Bekinschtein, and Chennu 2019; Canales-Johnson et al. 2020; Goupil and Bekinschtein 2012).

### Subjective midline modulated by alertness

Third, we used psychophysics to quantify the modulation of the subjective midline per participant by alertness levels. For this we fit a cumulative normal function (see methods section) to the proportion of rightward responses (per participant) under each stimulus condition from −60° to 60° from the midline under both, alert and drowsy periods. The mean of the function referred to as ‘bias’ is the subjective midline or spatial bias (where participants have 0.5 chance of pressing left or right responses) and the slope represents the sensitivity. Example fits of individual participants are shown in Figure 2C. Most participants (Figure 2D) had their bias point shifted to the left (as they became drowsy), indicating more left errors (overestimating the right side of space). A small proportion of participants had bias points shifted to the right. Overall, a paired sample t-test t(18) = 3.53, p<0.01, revealed a reliable difference in bias points between alert and drowsy periods. Next, we performed a paired sample t-test t(18) = −5.52, p<0.001 which revealed a reliable difference in slope between alert and drowsy periods. Further, we also plotted (Figure 2E) the mean of the psychometric fits of individual participants to show that the overall subjective midline has shifted to the left at the group level.

### Signal detection parameters modulated by alertness

Fourth, we used signal detection theory to understand the factors that modulate decision making under varying levels of alertness (Figure 2F). d′(sensitivity) showed extreme evidence in favour of the alternative hypothesis (being modulated by alertness) with a Bayes factor (BF) = 101100 while criterion (response bias) only showed anecdotal evidence in favour of the null hypothesis (being not modulated by alertness) with BF = 1.17. This suggests that internal representations in the brain in terms of sensory/perceptual and noise distributions are modulated by alertness levels. Further, no evidence for response bias also suggests that participants were not arbitrarily pressing right responses for uncertain stimuli with drowsiness.

To summarise, the behavioural results hint that the first two stages of perceptual and evidence accumulation processes are affected by decreasing alertness and that the final stage of motor implementation (response bias/criterion) may be less affected.

### Alertness modulates sequential sampling model parameters of evidence accumulation

Next, we aimed to quantify the different elements of the decision-making process using drift-diffusion modelling. The drift-diffusion model captures the optimal procedure involved in performing a 2-alternative forced choice (2AFC) task under sequential sampling framework. It assumes that the observer accumulates evidence for one or other alternative in every time step (Ratcliff et al. 2016), until integrated evidence reaches a threshold to decision (Figure 3A). The localization of tones to the left and right side of space is essentially a 2-choice task with the participant always forced to make a decision on the location of the tone. The model was implemented with a hierarchical Bayesian procedure using hierarchical drift diffusion model (HDDM) (See methods section). For the HDDMs, we fit the response of each participant instead of accuracy. This procedure is referred to as stimulus-coding, allows for the testing of several decision making parameters, and is critical to uncover different sources of bias (de Gee et al. 2017).

**Figure 3:**
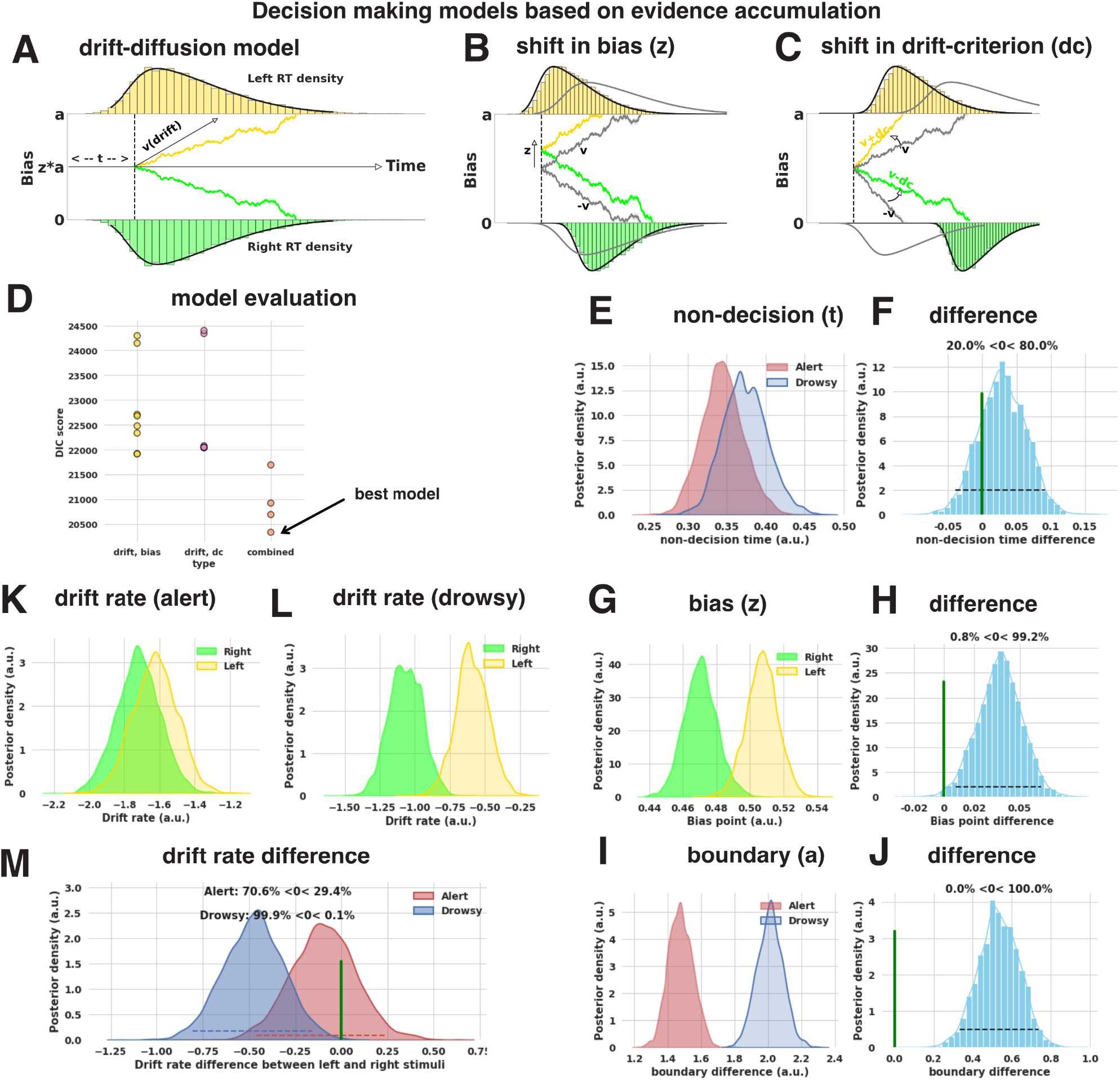
Evidence accumulation models: A) drift-diffusion model accounts for the reaction time distributions of responses across left and right stimuli (stimulus coding). ‘v(drift)’ indicates evidence accumulation rate, ‘a’ indicates the boundary separation, ‘z’ indicates the bias point, usually z = 0.5 for unbiased responses, ‘t’ indicates non-decision time composed of stimulus encoding and response execution. B,C) changes in response distributions can be explained by different sources of bias. Shift in starting point (z) or constant offset added to drift rate (drift criterion). grey lines indicate unbiased condition. D) In the first step, the appropriate source of bias (either z or dc) was identified. In our case, the source of bias was identified as ‘z’ based on the lowest DIC. Further, this bias (‘z’) was combined with different variations in ‘t’, ‘a’, ‘v’ and the best model with the lowest DIC was identified (refer to the methods section). E,F) posterior densities of ‘t’ in the best model and their differences indicate there is no reliable change (based on proportion of posterior overlap, here 20.0%) across alert and drowsy periods. G,H) posterior densities of ‘z’ in the best model and their differences indicate there is a reliable change (based on proportion of posterior overlap, here 0.8%) across left and right stimuli. I,J) posterior densities of ‘a’ in the best model and their differences indicate there is a reliable change (based on proportion of posterior overlap, here 0%) across alert and drowsy periods. K,L) posterior densities of ‘v’ for left and right stimuli across alert and drowsy periods in the best model. M) differences in the posterior densities of changes in drift rate indicate strong evidence (overlap reduces to 0.1% from 29.4%) in favour of change in drift rate across stimuli in the drowsy period compared to alert periods.

We examined models with different sources of bias: starting point - z (Figure 3B) or constant slope - dc (Figure 3C) and identified the best model as one with the lowest DIC (see methods section). The best model showed that drift-rate (v) varied according to state (alert or drowsy) and stimulus (left or right), bias-point (z) varied according to stimulus (left or right) but not state, non-decision time (t) varied according to state (alert or drowsy), and decision threshold (a) varied according to state (alert or drowsy). This furthers the claim that the changes in alertness place a higher burden in the evidence accumulation process (drift rate), and less on the motor implementation or stimulus encoding of the task (non-decision time). The best model was then analysed for differences in posterior densities of parameters. For this purpose, we used a Bayesian estimate which is more informative and avoids the arbitrary choices of significance level implemented for specific statistical tests used by the frequentist based methods (Kruschke 2012).

In the best model, the proportion of posterior overlap in the non-decision time (between alert and drowsy states) was 20.0% (Figure 3F). This indicates that the non-decision time was not reliably different between the different states. Next, the proportion of posterior overlap in the bias point (between left and right stimuli) was 0.8% (Figure 3H). This indicates that the bias-point was reliably different between the different stimuli. Next, the proportion of posterior overlap in the boundary (between alert and drowsy states) was 0% (Figure 3J). This indicates that the boundary was reliably different between the different states. Importantly, only the drift rate varied with respect to both the stimuli and the state, suggesting that the difference in evidence accumulation (indicated by the drift rate) is responsible for the difference in error rates between left and right stimuli in the low alertness state, as shown in the previous analyses. Here, the proportion of posterior overlap (between left and right stimuli) in the drift rate for alert trials was 29.4% and reduced to 0.1% for drowsy trials (Figure 3M). Thus the drift-rate (evidence accumulation rate) was reliably different between left and right stimuli on drowsy trials in comparison to the alert trials.

To summarise, the behaviour modeling results hint that only the central evidence accumulation process (indicated by drift rate) are affected by decreasing alertness, whereas processes responsible for perceptual encoding and response execution may not be modulated.

### Spatial and temporal signatures involved in spatial localization across alert and drowsy periods

Here, we were interested in identifying the neural signatures involved in the performance of this task. For this purpose, decoding involves identifying the stimulus (Y - left or right tone) presented from the EEG data (X). This process involves the identification of the W (classifier weights) that can produce the transformation, Y_t_ = W_t_X_t_ where ‘t’ represents time (Figure 4A,B). The performance of the classifier (W) is evaluated by training and testing the data at each time-point (t) using area under the curve (AUC) as measure. In Figure 4C, the shaded region represents those periods reliably decoded (p<0.05) (see methods for more details).

**Figure 4:**
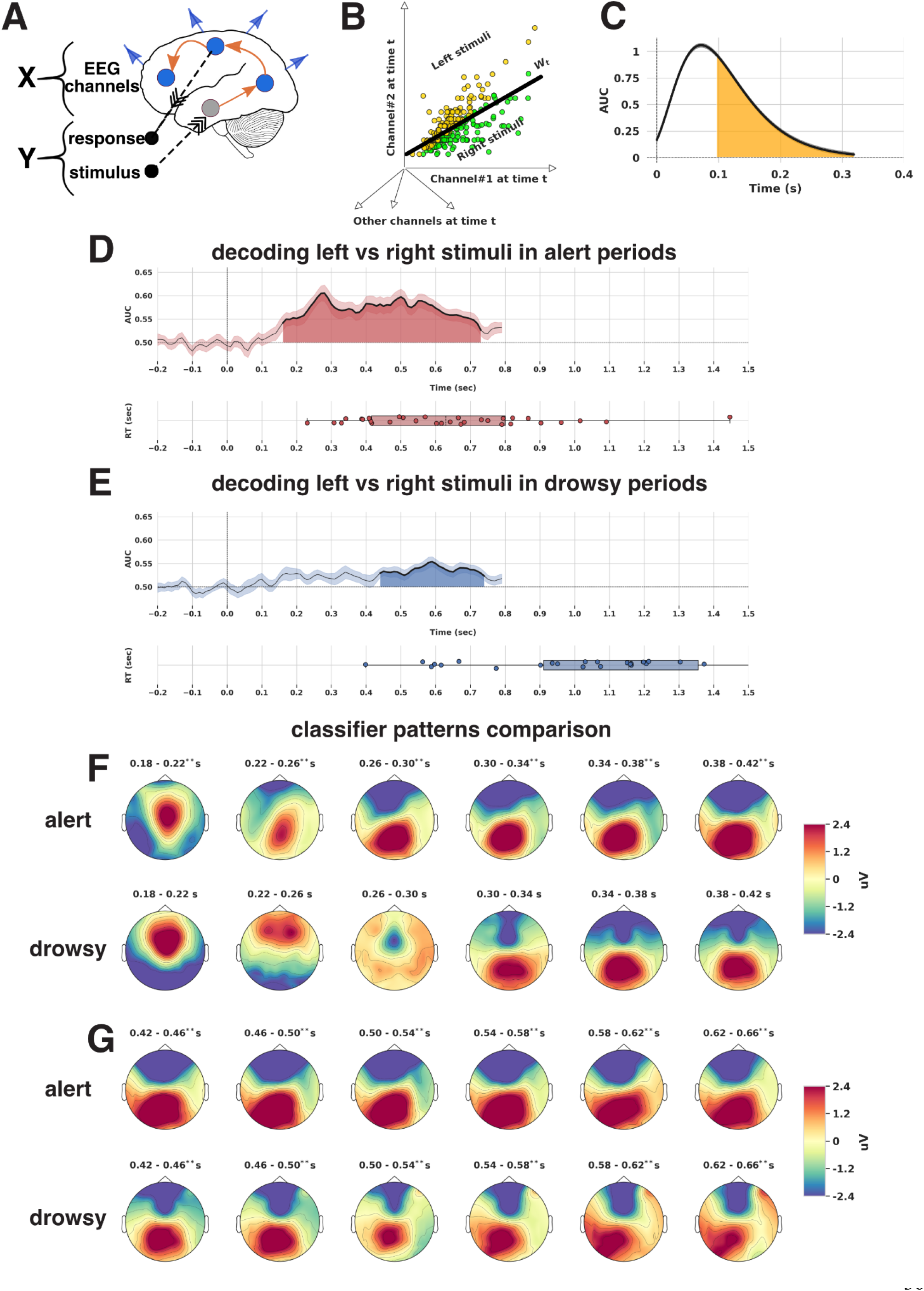
Temporal decoding (stimuli): A) decoding consists of identifying Y (stimuli) from X (EEG). The model thus consists of Y_t_ = W_t_X_t_ , where ‘t’ represents time and W_t_ represents transformation (classifier weights). Here, the predicted output Y consists of categorical labels (left/right tone) predicted from brain activity (X) B) classifier weights W_t_ are determined by the optimal separation of different classes (here left, right stimuli). C) decoding performance is assessed using the area under curve (AUC) where shaded regions represent reliably different time periods (p<0.05). D,E) AUC under alert and drowsy periods, where the classifier was trained to discriminate between targets of left and right stimuli. Shaded regions show time periods with reliable (p<0.05) discriminatory power, computed with a cluster based permutation test. The mean reaction times (RT) of participants are plotted below. F) Comparison of coefficients of classifier patterns in the early time periods. The early time periods highlight topographical differences in the frontal, posterior parietal and central electrodes. G) Comparison of coefficients of classifier patterns in the later time periods. The later time periods highlight topographical differences in the frontal and central electrodes. ** indicates reliable discriminatory in the corresponding period in temporal decoding.

We found that when participants were alert the reliable decoding of stimuli (Figure 4D) started at 160 ms after the stimulus and lasted until 730 ms (cluster permutation, p<0.05) with mean AUC of The peak discriminatory power was at 280 ms (AUC = 0.60). The average AUC between 200-300 ms was 0.58 ± 0.007, p<0.05, between 300-400 ms was 0.57 ± 0.002, p<0.05. However when the participants became ‘drowsy’ the decoding of stimuli (Figure 4E) shifted to 440 ms after the stimulus was presented and lasted until 740 ms (reliable with cluster permutation, p<0.05) with mean AUC of 0.53. The peak discriminatory power was at 590 ms (AUC = 0.55). The average AUC between 440-500 ms was M = 0.53 ± 0.001, p<0.05, between 500-590 ms was M = 0.54 ± 0.003, p<0.05. This suggest that the processes related to the discrimination between left and right stimuli under drowsy conditions may cease to be directly informative to the decision in the early processes of sensory encoding and would only start later, around ~400ms, to show neural differentiation.

Further to this, the lower discriminatory power points to a potentially less efficient (lower decodability, longer time duration, higher variability) process of central evidence accumulation, or to a different neural implementation of the processes during low alertness.

Next, we plotted the coefficients of the classifier patterns (derived from classifier weights W) in the scalp electrode space for further neurophysiological interpretation. We decided to compare the classifier patterns across early (<300ms), and later (>400ms) time periods. To compare patterns in the early time periods across alert and drowsy periods, we plotted the same for every 40 ms between 180 ms to 420 ms in Figure 4F. For the alert periods, the pattern between 180-220 ms indicates a strong involvement of signal in the fronto-central electrodes, whereas in the corresponding time periods under drowsy, the pattern seems to have shifted to more frontal electrodes although its contribution may be minimal as it was not reliably decodable. Under alert periods, the pattern shifts to more posterior regions (centro-parietal electrodes in the right side of the scalp) between 220 ms to 260 ms whereas under drowsy the pattern stays in the frontal electrodes itself, still not reliably decodable in the unconstrained voltage decoding. Further from 260-300 ms the pattern shifts to more parietal and occipital sites under alert periods and is only weakly parietal in the drowsy periods.

The comparison of the classifier patterns across alert and drowsy periods (in early time periods), reveals different topographies that point to a clear differential processing of information that would further map onto shifted perceptual and central evidence accumulation stages of cognitive processing (Sigman and Dehaene 2005) possible affecting the central accumulation. To compare patterns in the later time periods across ‘alert’ and ‘drowsy’ periods, we plotted the same for every 40 ms between 420 ms to 660 ms in Figure 4G. Particularly, the patterns in the alert periods have a lower frontal activity and higher posterior activity, whereas in the drowsy periods, the activity in both the frontal and posterior sites is much more localised although decodability is overall lower. Although only suggestive, this change in the decodability intensity and distribution suggest a delayed processing dynamics and or a change in the way of processing as alertness decreases.

The descriptive analysis of the classifier patterns indicate differences between alert and drowsy periods. To establish the spatial and temporal signatures of such differences we performed a cluster permutation test and identified regions where activity patterns in alert periods are different from drowsy periods. This analysis resulted in the identification for four clusters (Figure 5).

a. Cluster #1 (alert activity > drowsy activity): indicates early periods (150 ms to 330 ms), where activity is concentrated in parietal, central, posterior electrode sites. These spatial patterns are likely to be involved in early perceptual/evidence accumulation processes during the alert periods, as the majority of responses only start occurring from 400 ms onwards.
b. Cluster #2 (alert activity > drowsy activity): indicates later periods (430 ms to 650 ms), where activity is concentrated in occipital, parietal, central, posterior electrode sites. These spatial patterns are difficult to interpret as the reaction times overlap with the corresponding time periods. Hence this decoding pattern could be an amalgamation between central evidence accumulation and motor implementation processes.
c. Cluster #3 (drowsy activity > alert activity): indicates early periods (170 ms to 360 ms), where activity is concentrated in frontal electrode sites. These spatial patterns are likely to be involved in early perceptual/sensory encoding processes during the drowsy periods, as the majority of responses only start occurring from 900 ms onwards. The temporal neurodynamics show delayed patterns in drowsy compared to awake.
d. Cluster #4 (drowsy activity > alert activity): indicates later periods (380 ms to 650 ms), where activity is concentrated in left frontal electrode sites. These spatial patterns are likely to be involved in central evidence accumulation related processes. However it is difficult to interpret as the corresponding time periods during alert mildly overlap with response related activity.

**Figure 5:**
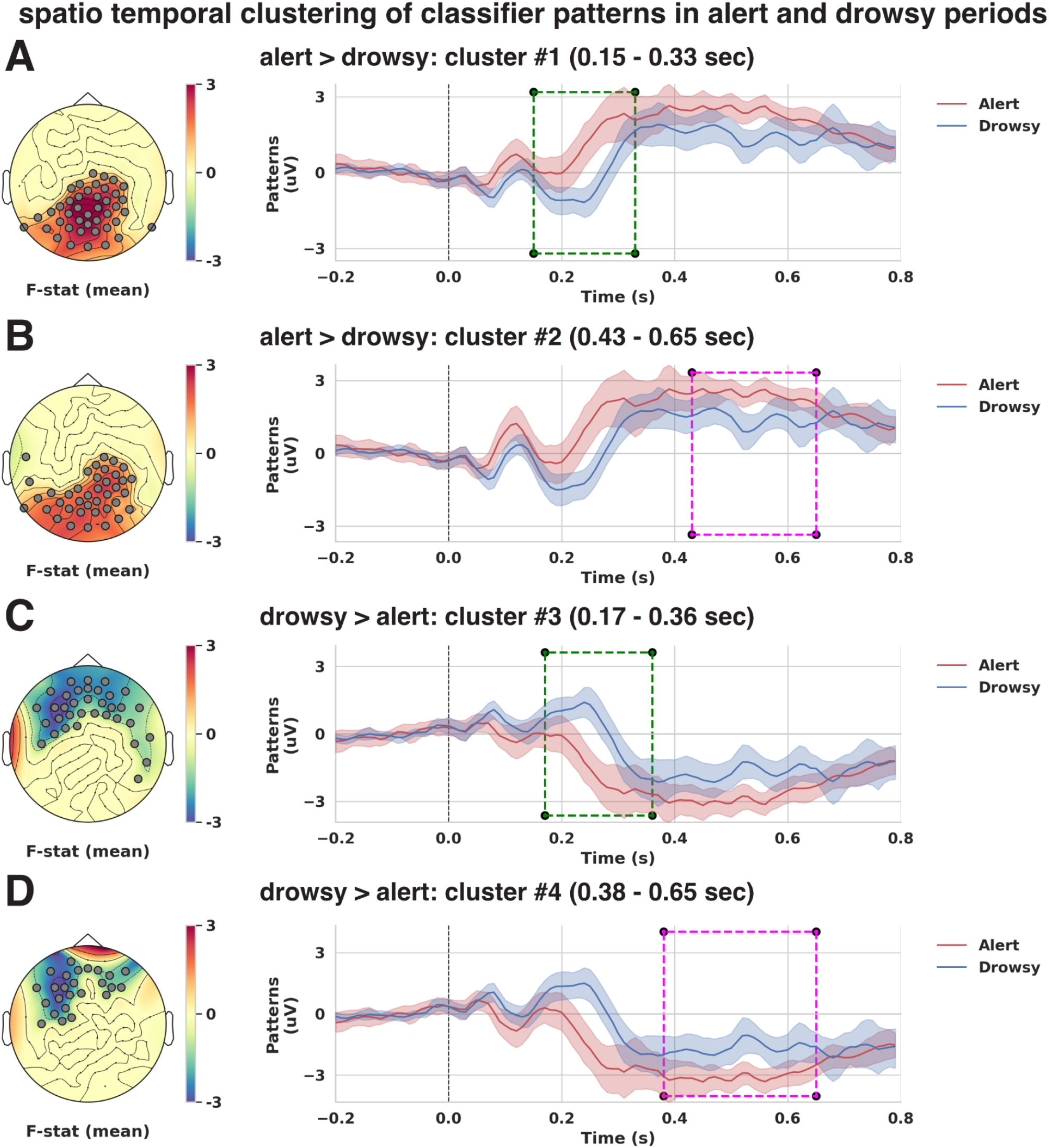
Classifier pattern differences: To identify differences between the classifier patterns in Figure 4 (F,G) we performed spatio-temporal clustering using a permutation test. A) Cluster#1 indicates early periods (150 - 330 ms after stimulus) where alert activity is higher than drowsy periods. B) Cluster#2 indicates later periods (430 - 650 ms after stimulus) where alert activity is higher than drowsy periods. In terms of spatial locations, the early and later clusters differ in right parietal sites, central, middle electrode sites. C) Cluster#3 indicates early periods (170 - 360 ms after stimulus) where drowsy activity is higher than alert periods. D) Cluster#4 indicates later periods (380 - 650 ms after stimulus) where drowsy activity is higher than alert periods. In terms of spatial locations, the early and later clusters differ in left frontal sites, middle electrode sites.

To summarise, these results indicate differences in spatial patterns related to early sensory/evidence accumulation related process (cluster #1,3) across alert and drowsy periods. The next step is to identify and tease apart motor preparation related processes occurring during later time periods.

### Spatial and temporal signatures of motor implementation across alert and drowsy periods

In classic cognitive processing model frameworks, motor implementation occurs after the central evidence processes, when information of the decision is routed to the premotor network, this is assumed to be mostly a serial process. We preprocessed the EEG data now locked to responses (see methods section) and decoded the response hand itself (irrespective of the stimulus presented). These response related patterns thus would depict information related to the motor implementation process irrespective of the decision related processes.

The decoding performance: AUC (Figure 6A,B) starts ramping up from −210ms (alert), −150ms (drowsy) from the response time, and increases when the button is actually pressed at 0 sec, reaching the peak soon after. In both cases, awake and drowsy, the neural decoding results last till 190ms. These results are in line with motor implementation related processes commonly reported around 200 ms before the actual motor response, gradually ramping up to produce the response itself. It is also interesting to note that the AUC for alert is always higher than drowsy, indicating less variability in neural concerted activity, leading to better classifier performance without time span differences, this may indicate a more efficient motor implementation in the awake state.

**Figure 6:**
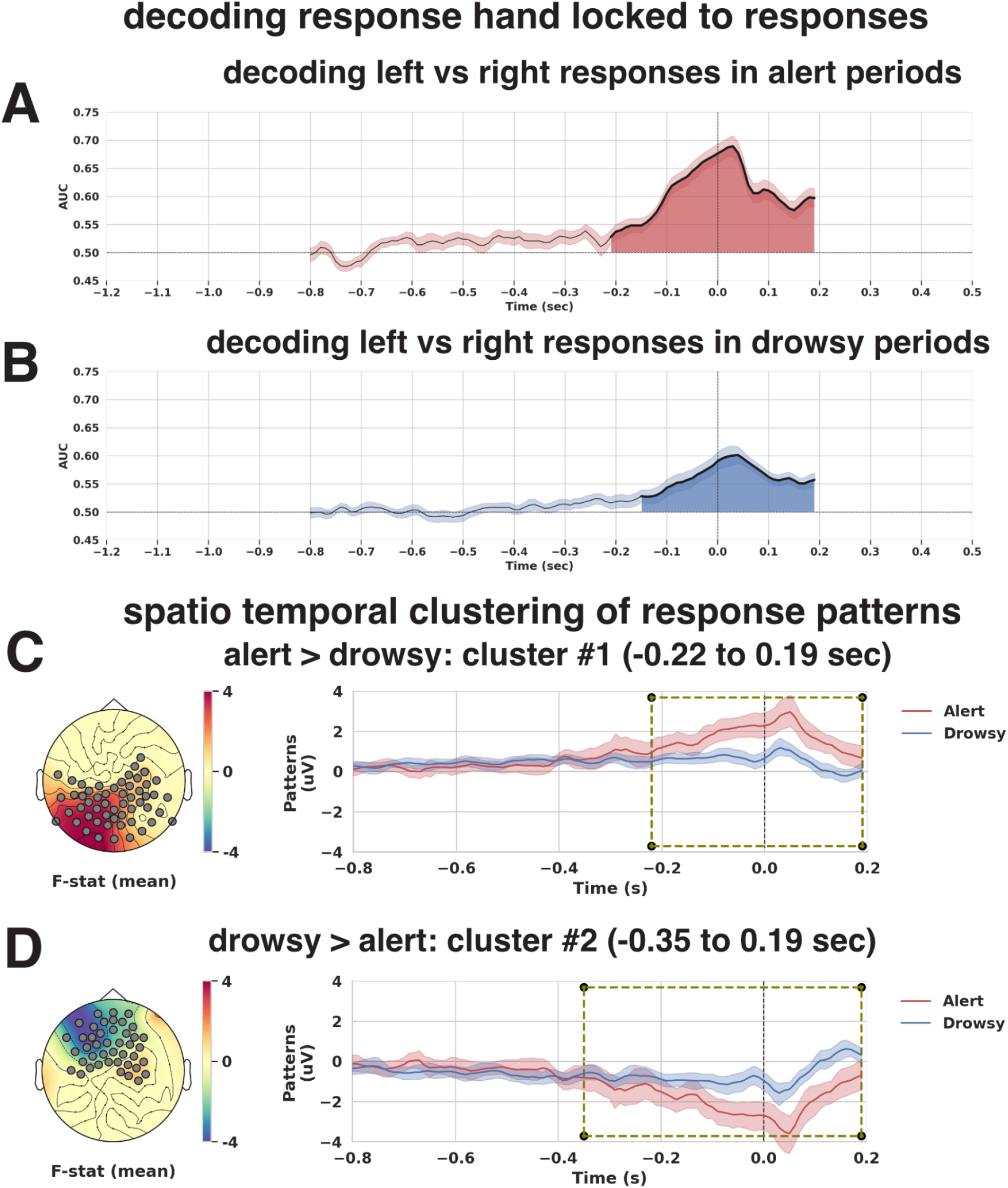
Temporal decoding (responses): To identify differences in motor preparation (response related), we decoded the left and right button press events irrespective of stimuli presented. A,B) AUC under alert and drowsy periods, where the classifier was trained to discriminate between targets of left and right button presses. Shaded regions show time periods with reliable (p<0.05) discriminatory power. The time point of 0 corresponds to the actual button press event. The ramp up of the AUC under both conditions starting from 210ms (alert), 150ms (drowsy) before the actual response indicates the ability of the decoder to detect neural signatures related to responses. Spatio-temporal clustering of differences in classifier patterns produces two clusters with similar time periods in contrast to the clusters (with early and later time periods) produced by stimuli related classifier patterns. C) Cluster#1 indicates periods (−220 to 190 ms) where alert activity is higher than drowsy periods. D) Cluster#2 indicates periods (−350 to 190 ms) where drowsy activity is higher than alert periods. In terms of spatial locations, the clusters differ in frontal, central, middle, parietal electrode sites. The presence of a single cluster with similar time periods across alert and drowsy periods indicate that motor related processes start appearing at similar time periods and hence do not change with modulations in alertness levels.

To establish the spatial and temporal signatures of differences in motor related processes we performed a cluster permutation test of classifier patterns and identified regions where activity patterns in alert periods are different from drowsy periods. This analysis resulted in the identification of two clusters (Figure 6C,D).

a. Cluster #1 (alert activity > drowsy activity): indicates periods (−220 ms to 190 ms), where activity is concentrated in premotor, parietal, posterior electrode sites.
b. Cluster #2 (drowsy activity > alert activity): indicates periods (−350 ms to 190 ms), where activity is concentrated in premotor, frontal electrode sites.

The existence of a single cluster (in time) across both alert and drowsy periods further indicates that motor related processes do not necessarily vary in time with modulations in alertness levels. Furthermore for both alert and drowsy periods the start of the decoding happens during similar time periods around −200ms and lasts till similar time periods ~190ms.

In summary, these results indicate that motor preparation related processes are similar during both alert and drowsy trials, suggesting this process may not be critical to the changes in decision making in low alertness when compared to evidence accumulation related processes. Although the efficiency of the implementation may be lower in drowsy this could be due to differences in signal to noise with alertness or to changes in the neural implementation of the action itself.

The above analysis related to decoding of stimuli and responses indicate that most likely only central evidence accumulation related processes are modulated by alertness. However the decoding analysis in the late clusters could be confounded by response related processes occurring during alert periods. Hence we devised the next step of the analysis where we use parameters derived from the drift diffusion modelling to understand how different elements of the evidence accumulation are represented in the neural domain. This analysis is determined by the drift diffusion model as a mathematical implementation of decision making theory, and it guides neural exploration to capture the brain dynamics echoes of the evidence accumulation process.

### Neural signatures of evidence accumulation of decision making modulated by alertness

To take advantage of the information gained by the drift diffusion modelling we decided to capture the neural implementation of the evidence accumulation parameters in the decision making process, and how it might differ in wakefulness and low alertness periods. We specifically aimed at capturing the change in the neurobehavioural dynamics of the evidence accumulation process in the awake and drowsy periods. To establish this we developed a novel method based on the drift-diffusion analysis shown earlier. We used the best model (from the drift diffusion analysis earlier) and computed trial-by-trial influence of a brain covariate (ERP) on the drift diffusion model parameters (see methods section). Further we compute the proportion of the overlap of the posterior distributions of the trace obtained for both left and right stimuli separately under alert and drowsy conditions (Figure 7A). This analysis is repeated for each participant and we obtain regression patterns, similar to the classifier patterns but guided by the drift rate, per participant.

**Figure 7:**
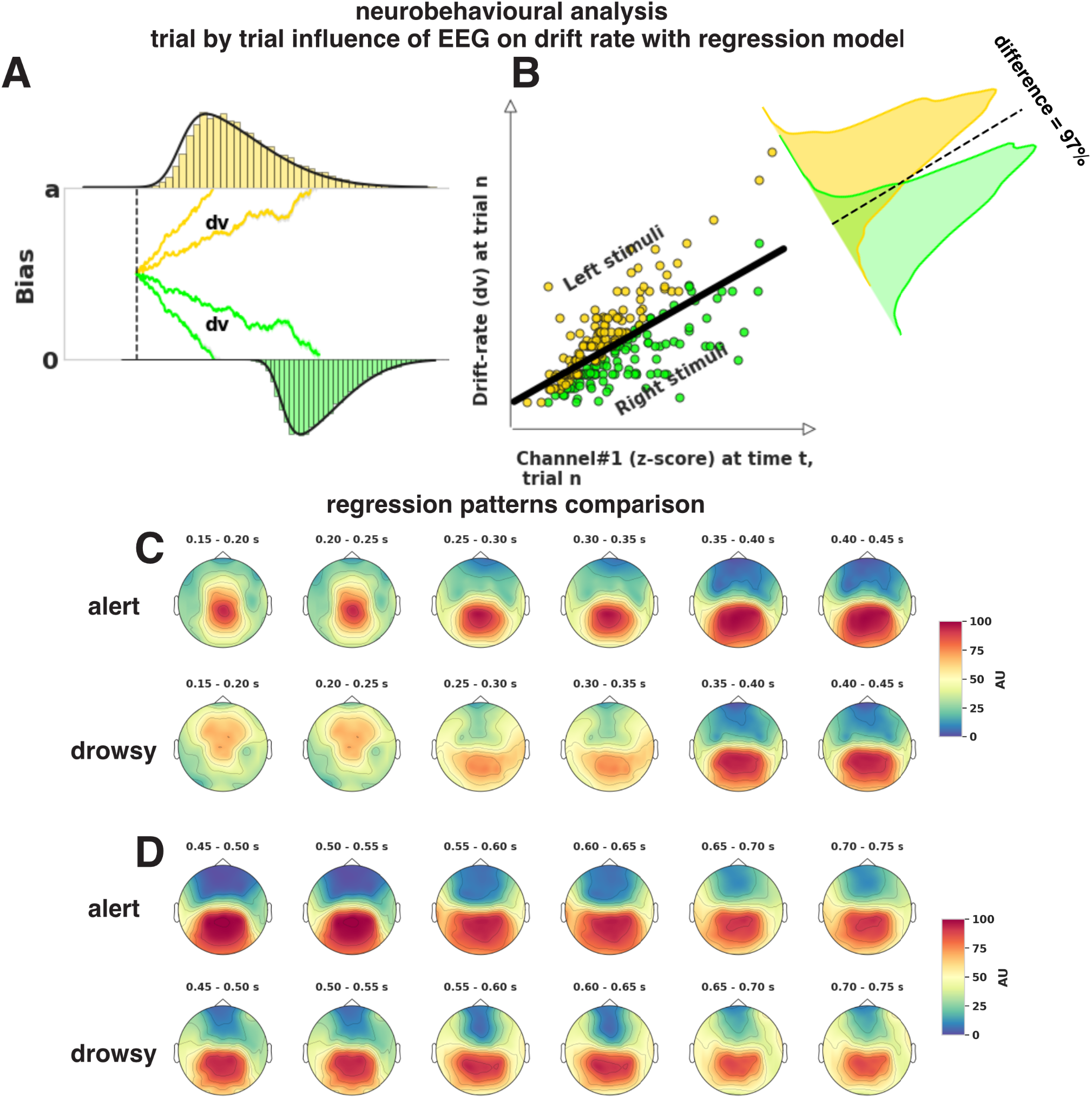
Regression patterns: A,B) EEG data (z-scored and averaged per 50 ms) was regressed against drift rate (v), the difference in the posterior of regression rate was quantified per electrode per time-point per participant. C,D) mean regression patterns in alert and drowsy periods projected in electrode space show a classic centro frontal to parietal progression from 150ms to 750ms. The early time periods highlight topographical differences in the frontal, middle, central parietal electrodes.

Next, we plot the regression patterns in alert and drowsy periods. The patterns in the alert periods (Figure 7C,D) indicate that the topography focuses initially in the fronto-central electrodes (150 ms to 250 ms), which then shifts to centro-parietal electrodes (250 ms to 350 ms), similar to the classifier analysis (decoding patterns from figures 4F and 4G). The patterns in the drowsy periods (Figure 7C,D) indicate that the topography initially focuses on the frontal electrodes (though weaker compared to alert periods) in the interval from 150 ms to 250 ms. Further the patterns shift to more central electrodes (again weaker when compared to alert periods) in the interval from 250 ms to 350 ms.

The descriptive analysis of the regression patterns indicate differences between alert and drowsy periods. To establish the spatial and temporal signatures of such differences we performed a cluster permutation test and identified regions where activity patterns in alert periods are different from drowsy periods. This analysis resulted in the identification of two clusters (Figure 8).

a. Cluster #1 (alert activity > drowsy activity): indicates early periods (125 ms to 275 ms), where activity is concentrated in parietal and central electrode sites. These spatial patterns are likely to be involved in early perceptual and evidence accumulation processes.
b. Cluster #2 (drowsy activity > alert activity): indicates early periods (175 ms to 325 ms), where activity is concentrated in frontal/central electrode sites. These spatial patterns are likely to be involved in early perceptual/sensory encoding processes during the drowsy periods.

**Figure 8:**
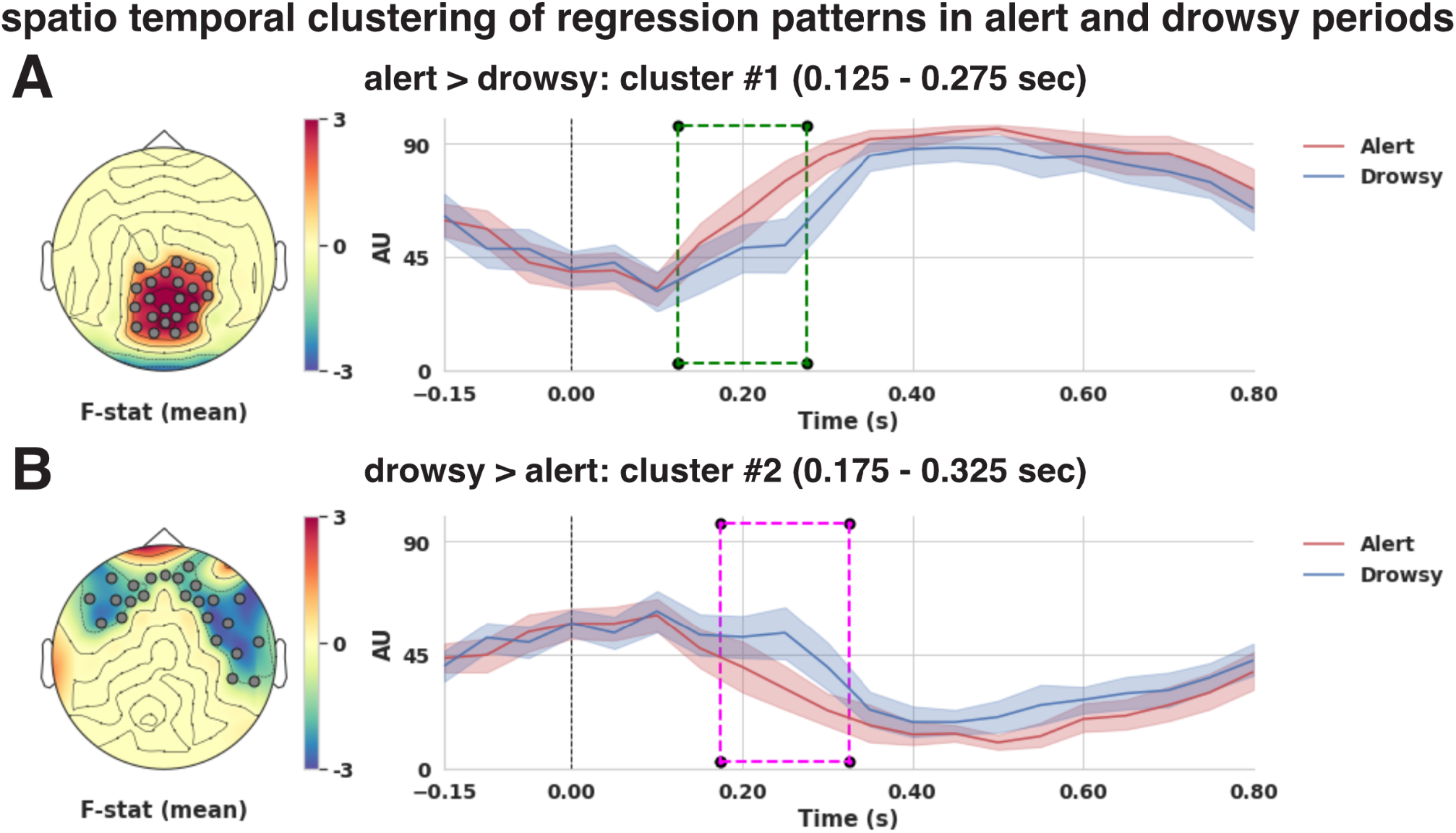
Regression pattern differences: spatio-temporal clustering of differences in regression patterns shown in Figure 7 (C,D) between alert and drowsy periods. A) Cluster#1 indicates early periods (125 - 275 ms after stimulus) where alert activity is higher than drowsy periods. In terms of spatial locations, the cluster#1 identifies electrodes in right parietal sites, central, middle electrode sites. B) Cluster#2 indicates early periods (175 - 325 ms after stimulus) where drowsy activity is higher than alert periods. In terms of spatial locations, the cluster #2 identifies electrodes in left/right frontal sites, right middle electrode sites.

These analyses indicate spatial/temporal regions where the alert and drowsy periods differ in terms of sensory encoding and central evidence accumulation related processes. Furthermore, we have also teased apart these differences from motor related processes by directly using a parameter (drift-rate) from the hierarchical drift diffusion modelling.

### Alertness modulates frontoparietal cortical neural patterns in perceptual decision-making

Next we performed an exploratory analysis to identify the putative neural sources of these differences (Figure 9) using a source reconstruction procedure to project the classifier patterns of the different clusters back to their neural sources separately. In the source space, we performed cluster permutation tests to identify where in the cortex the activity in alert periods is higher than in drowsy periods. We used the actual value of the source activity (instead of absolute values), as we are interested in the distance between the two different patterns and not in overall activity levels.

a. Cluster #1: This cluster corresponds to regions where activity in alert periods is higher than drowsy periods. These regions are putative locations for early sensory, evidence accumulation related processes in alert periods. The analysis reveals regions (inferior parietal lobule) previously associated with the VAN - ventral attention network (Yeo et al. 2011; Corbetta and Shulman 2011) in the right hemisphere. Further reveals regions (superior parietal) that are part of the DAN - dorsal attention network (Yeo et al. 2011; Corbetta and Shulman 2011) and central processing of flexible information (Fedorenko, Duncan, and Kanwisher 2013).
b. Cluster #2: This cluster corresponds to regions where activity in drowsy periods is higher than alert periods. These regions are putative locations for early sensory related processes in drowsy periods. The analysis reveals regions (superior temporal gyrus, inferior frontal gyrus, supramarginal gyrus, transverse temporal, temporoparietal, fronto-parietal - middle frontal gyrus) regions in the right hemisphere also associated with the ventral attention network (Yeo et al. 2011; Corbetta and Shulman 2011).

**Figure 9:**
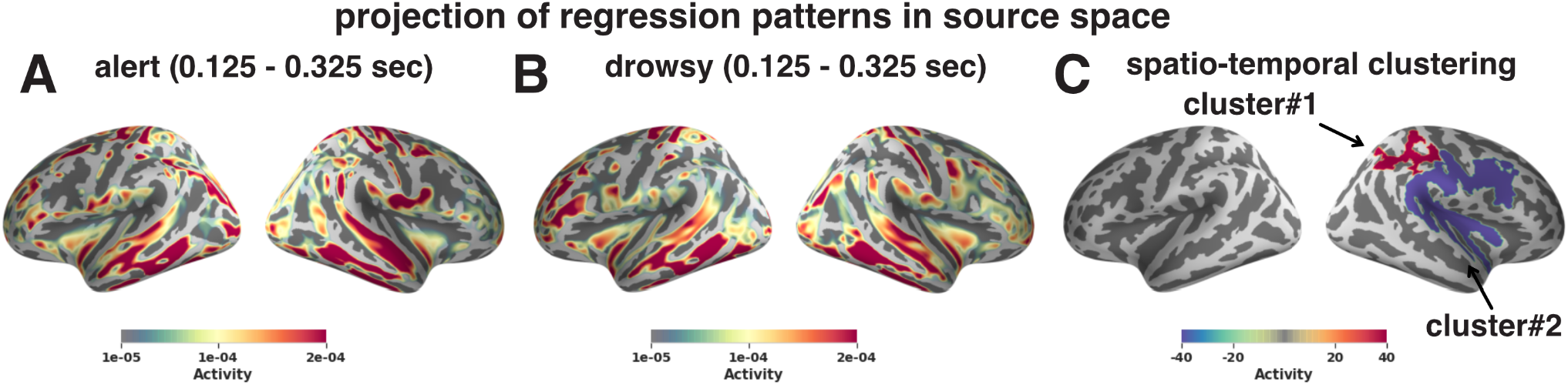
Regression patterns projected into source space: To identify the putative regions/networks in the brain related to perceptual decision-making and its modulation by alertness we projected the regression patterns in Fig 8 to source space. A) average regression patterns in the alert periods (125 - 325 ms) projected into source space. B) average regression patterns in the drowsy periods (125 - 325 ms) projected into source space. C) cluster#1 (alert > drowsy) identifies differences in regions related to the dorsal and ventral attention network in the right superior and inferior parietal regions respectively. cluster#2 (drowsy > alert) reveals differences in regions related to the ventral attention network in the superior-temporal, fronto-parietal regions.

To summarise, these results indicate that during alert periods: the early sensory/central accumulation related process (until 300ms from stimulus presentation) are located in the regions corresponding to the dorsal attention network (which is specialised for spatial attention). For the drowsy periods: the early sensory encoding related process (until 300ms from stimulus presentation) are located in the regions corresponding to the fronto-parietal, ventral attention network, further during later periods (after 300ms) evidence accumulation related process are located in the regions corresponding to fronto-parietal network. This indicates that fronto-parietal regions of the brain are further recruited for the late decision-making processes implemented in the drowsy periods, pointing to a cortical network reconfiguration in the evidence accumulation between normal and low alertness periods.

## Discussion

In this study we characterize cognitive and neural dynamics of perceptual decision making while participants are falling asleep. First we established, using multi-level modelling, that errors for left-sided stimuli (compared to right) increased as participants become drowsy, in convergence with (Bareham et al. 2014) but with a different study design, analysis methods and participants sample. Further, we fitted the proportion of rightward responses of individuals and identified the subjective midline, showing that shifts to the left (more rightward responses) as people become drowsy. These echoes line-bisection studies (Jewell and McCourt 2000) showing that healthy individuals bisect lines to the left of the veridical midline, and further supports meta-analysis studies suggesting rightward shift in attention with lower levels of alertness (Chandrakumar et al. 2019). Additional studies (Benwell, Harvey, and Thut 2014) have also identified possible neural origins of such bias as task independent activity in the ventral attention network, however, they did not examine whether trial to trial alertness of individuals was modulated and how the different aspects of the decision making process were affected. Several studies (Corbetta and Shulman 2011) have shown that activity in the ventral attention network interacts with the alertness system, and that low alertness, as shown directly here, could be the root cause.

Next, using signal detection theory we found that sensitivity (d′) had strong evidence of modulation by low alertness whereas criterion showed anecdotal evidence in favour of no modulation, complementing early deprivation studies in vigilance tasks (Deaton, Tobias, and Wilkinson 1971). This suggests that the sensory/perceptual representations have indeed been modulated by alertness, and that participants were not arbitrarily pressing more rightward responses (in the face of uncertain stimuli) as alertness decreases. Further, using the sequential sampling model framework, we showed that in the best model, only the drift-rate was reliably modulated by both alertness and stimuli, whereas starting-point was only affected by stimuli (reliable), and boundary (reliable) and non-decision time (not-reliable) were affected by alertness. This shows that alertness indeed differentially modulates evidence coming from the left side of space compared to the right. Decision-making studies in the visual modality (Smith and Ratcliff 2009) have shown that drift rates are closely related to the strength of stimulus encoding, partially affected by attentional strength and evidence accumulation to a decision threshold. Further studies have also shown neural correlates of such parameters in the EEG (Nunez, Vandekerckhove, and Srinivasan 2017; O’Connell, Dockree, and Kelly 2012) and its relationship with stimuli strength. Hence we interpret the effect of alertness modulation on drift rate as acting specifically on spatial attention affecting the stimulus encoding, as well as in central processes of evidence accumulation. Finally, it is important to compare the parameters of the signal detection theory (SDT) with the drift diffusion model (DDM) (Jun et al. 2021). The changes in sensitivity (d′) in SDT could be directly related to the changes in the boundary parameter (a) and drift rate (v) in the DDM. Here the changes in d′ are in agreement with both ‘v’ and ‘a’ as both are shown to be modulated by alertness levels. The changes in criterion (c) in the SDT correspond to sources of bias and they could be originating from the stimulus: corresponding to drift criterion (dc) in the DDM or originating from the response: corresponding to starting point (z) in the DDM. With the evaluation of competing models we show that models with ‘z’ outperforms the models with ‘dc’, further ‘z’ varies only stimulus direction (left or right) and is not modulated by alertness. This agrees with previous studies showing an initial spatial bias (Benwell et al. 2013; Learmonth et al. 2015) even before any experimental modulation (like ‘time on task’ or ‘alertness’). Here the evidence for the modulation of ‘z’ in the DDM across stimuli direction and not alertness levels agrees with the non-modulation of criterion in the SDT across alertness levels.

Next using time-multivariate decoding, we showed that under alert conditions the decoding started early, at 160 ms and lasted until 730 ms, however, under drowsy conditions this was delayed to 440 ms and lasted until 740 ms. This ~280 ms delay may indicate that, under lower alertness, the brain requires a longer time to decipher -or process- the direction of stimuli as suggested by slower reaction times and a change in the drift rate, or alternatively, that the neural code in this window is too noisy to be captured by this neural decoding. Furthermore, the overall strength of decoding was systematically higher in alert than drowsy, suggesting that the brain processes responsible for decision making are less variable under the alert compared to the drowsy condition, or that there is less efficient decision processing during low alertness (see (Killgore 2010) for an account on sleep deprivation on cognition).

The different neural time clusters potentially point to a first stage of information encoding and second of central evidence accumulation (Kelly and O’Connell 2015) that others have mapped to perceptual/central decision processes (Sigman and Dehaene 2008). If that is the case, the first neural signal cluster in drowsy (cluster#3 in Figure 5C), showing changes in the spatial pattern and a lack of reliable decoding may be interpreted as less efficient encoding and transfer of information in low alertness or may be using different network dynamics that are not captured by the direct time-voltage decoding. The second cluster (cluster#4 in Figure 5D), more similar between states, may signal a lower efficiency of central evidence accumulation, as it happens around the time the central decision process occurs in cognitive control (Ho, Brown, and Serences 2009). Furthermore, the second window of processing may also show part of the motor implementation processes at least for the awake condition (cluster#2 in Figure 5B), where the responses overlap for several participants (Figure 4D). The overlapping processes from ~400ms in awake, a mixture of central evidence accumulation and motor implementation of the decision, depending on the participant, are easily dissociated when drowsy, when the motor plan is implemented ~500ms later (around 900ms). This is complemented by the response-locked analyses that seem to confirm a common motor implementation, and it further guides the subsequent analysis.

Further analyses of the classifier patterns suggested specific signatures of early perceptual and late central cognitive processes. In the alert condition, the early pattern (180-220 ms) indicates activity over frontocentral electrodes, further shifting to more central and parietal sites later on. In the drowsy condition, the pattern initially starts at frontal sites (though not strong enough to be decoded) and shifts to central and parietal sites, later on from 380 ms onwards. Thus the formation of central, parietal patterns (possibly related to the first stage of encoding, evidence accumulation) takes longer or starts later in drowsy periods compared to alert. The spatial and temporal distribution of this centro-parietal decoding pattern resembles both a classic P2 and early P3, which has been dubbed as the build-to-threshold signal (Twomey et al. 2015), and the centro-parietal positivity (CPP), a more specific signal of evidence accumulation (Loughnane et al. 2016) respectively. To disambiguate the neural implementation of the evidence accumulation, we developed a novel method using regression patterns that identified differences in drift rate across alert and drowsy periods and further tested neural differences between alertness conditions.

Cluster#1 (alert>drowsy) showed activity in inferior parietal source regions that resemble the patterns of the ventral attention network. Although we cannot directly differentiate between early perceptual encoding and central accumulation processes, this cluster most likely corresponds to both in the alert periods. This part of the process of the spatial decision shows the early involvement of regions that have been linked, in several studies (Dietz et al. 2014; Shulman et al. 2010), to lesions in the inferior parietal lobule resulting in egocentric neglect. Furthermore, lower activity in the ventral attention network was also reported for neglect patients suffering from deficits in arousal (Corbetta and Shulman 2011). This agrees with the higher activity observed in the regression patterns for the alert periods compared to the drowsy periods of lower alertness. Further, it also revealed regions in superior parietal that have been associated with the dorsal attention network (Dietz et al. 2014; Shulman et al. 2010). Again these regions most likely correspond to the central evidence accumulation that is directly involved in evaluation of the spatial attention information.

Cluster#2 (drowsy>alert) showed activity in the transverse temporal, superior temporal gyrus, temporoparietal junction, fronto-parietal regions (inferior frontal, middle frontal gyrus) in the right hemisphere that can also be considered part of the ventral attention network. Several studies (Heilman et al. 1987; Corbetta et al. 2005) have reported lesions in the right ventral frontal cortex in patients suffering from both arousal related deficits and neglect to the right side of space. The highlighting of these regions in our study indicates that they are different from the alert periods and their higher activity in the regression patterns points to a potential source of spatial neglect caused by alertness deficits, which is similar to that observed in patient and studies in healthy volunteers with causal manipulation (Paladini et al. 2017). This could be dissociated in further studies, either with lesion metaanalysis or with virtual lesions in normal volunteers as we recently suggested (Bareham et al. 2018). We think this cluster corresponds to a combination of sensory encoding and evidence accumulation in the drowsy periods, highlighting an early cortical reconfiguration of the evidence accumulation process.

When participants are fully alert we think the information is processed initially by the ventral attention network, followed by the dorsal (specialised in spatial attention). Whereas when participants are drowsy the ventral attention network, although disproportionately affected in the right hemisphere (Corbetta et al. 2005; Corbetta and Shulman 2011; Heilman et al. 1987), is still involved in sensory encoding and early evidence accumulation processes. This reconfiguration in the ventral attention network is further propagated in the next stage, the dorsal attention network. This second part of the decision closely tallies with the proposal by Corbetta and colleagues to account for spatial neglect found in stroke patients suffering from lesions in right-hemispheric regions. But further, this network shows temporal and spatially extended recruitment of the parietal and frontopolar cortices to compensate for the direct effects of lower alertness, and that reconfiguration of the brain networks to relatively maintain performance may be a common and expected mechanism (Canales-Johnson et al. 2020).

As we lose consciousness, the neural system responsible for decision making adapts to the internal challenge of decreasing arousal and shows its resilience, exerting homeostatic regulations at the cognitive level to maintain performance. The attention and wakefulness fluctuations experienced in humans (and characterised in other animals) are common during the day and they are not only dependent on the circadian and sleep-wake regulation pressure (Borbély et al. 2016), but also on genetic, epigenetic, environmental, and life history factors that shape the alertness aspects of attention as well as the alertness aspect of arousal (Mitchell 2020; T. A. Bekinschtein et al. 2009).

From a decision-making perspective, we have added a missing level of description by characterising possible mechanisms of resilience of the neurocognitive processes elicited by decreases in tonic alertness. When moving from fully alert to a lower alertness state the brain tries to recruit and expand evidence accumulation related processes to fronto-parietal regions instead of temporoparietal regions, consistent with an increase in cognitive demand proposed by the Multiple Demand system (Duncan 2010). We interpret that the neural reconfiguration occurring in the transition from fully awake to decreased alertness brings different neural dynamics and changes the nature of the noise in the brain, forcing the cognitive system to exchange information differently. We put forward that low alertness negatively impacts the efficiency of the evidence accumulation processes, and differentially impacts performance, and the response of the system is to compensate by extending its neural processes in time and space to attempt to maintain performance. These findings highlight the neural dynamics of decision making when the external world remains unaltered, physical evidence held constant, but the internal milieu fluctuates exerting a modulatory influence on the cognitive decision making systems that caused them to reconfigure to solve the task at hand.

Transition of consciousness in the near-awake to light-decrease of alertness is emerging as a model for internally caused interactions (Comsa, Bekinschtein, and Chennu 2019; Tagliazucchi and Laufs 2014; Song and Tagliazucchi 2020; Canales-Johnson et al. 2020; Noreika et al. 2020). With lesion and pharmacological challenges, neuropsychology and cognitive neuroscience have tried to define the necessary and sufficient brain networks, areas and dynamics of the brain to implement cognition, revealing compensation, reconfiguration and plasticity (Adolphs 2016; Valero-Cabré et al. 2017; Yeung, Tzvetkov, and Atanasov 2018). Semi-causal effect of the internal interference exerted by the relative independence of the arousal system on the cognitive process, paves the way for the use of wakefulness, alertness and arousal challenges in a principled manner, adding new tools for cognitive brain research. Cognitive neuroscience uses models, correlational and causal methods to reach consensus about the underlying mechanism of thought (Krakauer et al. 2017). Here, we have followed a theoretically motivated question about perceptual decision-making systems, using behavioural modelling to understand the system, but at the same time causally modulating the neural networks with alertness decreases to uncover the mechanisms of brain function.

## ACKNOWLEDGMENTS

This research was funded by the Gates Cambridge Scholarship awarded to S.R.J and the Wellcome Trust Biomedical Research Fellowship WT093811MA awarded to T.A.B. as well as small departmental funding to S.R.J. and T.A.B. We thank Annamaria Laudini and Dritan Nikolla for assistance with data collection, Andrés Canales-Johnson, Will Harrison, Valdas Noreika and other members of the Consciousness and Cognition Lab in Cambridge for their valuable comments and support. We also thank lavazza for the continuous support with delicious and stimulating coffee.

## Dataset availability

The raw dataset associated with this study are present here: https://doi.org/10.5281/zenodo.5655443

## AUTHOR CONTRIBUTIONS

Conceptualization: S.R.J and T.A.B.

Methodology: S.R.J.

Software: S.R.J.

Investigation: S.R.J, C.A.B. and T.A.B.

Formal Analysis: S.R.J.

Resources: S.R.J, C.A.B. and T.A.B.

Data Curation: S.R.J and C.A.B.

Writing – Original Draft: S.R.J. and T.A.B.

Writing – Review & Editing: S.R.J, C.A.B, and T.A.B.

Visualization: S.R.J.

Supervision: C.A.B. and T.A.B.

Project Administration: S.R.J, C.A.B. and T.A.B.

Funding Acquisition: S.R.J and T.A.B.

